# “Fuzzy specificity” shapes diazotroph diversity and composition in nodulating plants of the Southeastern USA

**DOI:** 10.64898/2026.06.16.732771

**Authors:** D.J. Pantinople, P.D. Giram, J. Doby, S. Ahmed, T. Singh, N.J. Engle-Wrye, C.M. Siniscalchi, H.R. Jordan, R.P. Guralnick, R.A. Folk

## Abstract

Nitrogen-fixing symbioses, particularly those occurring in root nodules, are among the most consequential mutualisms in natural and agricultural systems and represent a globally important source of bioavailable nitrogen. Despite their importance, patterns of diversity and composition among diazotrophic symbionts—and the processes structuring those patterns in natural systems—remain poorly resolved, with competing hypotheses emphasizing ecological, or phylogenetic constraints on host–symbiont associations. Here, using a broad survey of nodulating plants from the southeastern United States, we examine how diazotrophic symbiont communities vary across host plant phylogeny, habitat context, and geographic origin. We find that host phylogeny is the primary determinant of symbiont composition, outweighing effects of fine-scale taxonomic identity. Symbiont associations are therefore structured mainly at deeper phylogenetic levels, consistent with phylogenetically constrained, or “fuzzy,” host specificity. Likewise, nodule community diversity—potentially reflecting variation in host control over infection—differs primarily among higher-level clades rather than among closely related taxa. Habitat context also shapes nodule communities, but its influence is secondary and most evident in undisturbed environments. As well, nonnative legumes harbor distinct symbiont assemblages despite occupying similar habitats, whereas distantly related legume clades share symbionts across habitats, highlighting interactions among phylogeny, ecology, and geographic history. Overall, our results show that host phylogeny exerts the strongest influence on nodule microbial communities, likely reflecting evolutionary divergence in symbiotic function across major host lineages.

## INTRODUCTION

One of the most important symbioses in terrestrial systems involves nitrogen-fixing bacteria and their plant hosts. Nitrogen, the critical building block of DNA, RNA, and amino acids, as well as a major component of chlorophyll and other biomolecules, is not directly available to plants from the atmosphere and is thus the key limiting resource of primary productivity in most ecosystems (Falkowski, 1997). Diazotrophs (microbes that fix nitrogen) subsist either as free-living soil or water organisms or have more direct associations with plant roots and aboveground biomass where they live in the intercellular spaces and are capable of diazotrophy (Roper and Gupta, 2016; Gupta et al., 2019; Hurek et al., 2002). Some plants directly recruit bacterial diazotrophs that can directly fix atmospheric nitrogen into its biologically available form, creating the basis of a remarkably successful symbiosis in natural and agricultural systems. The most prevalent and likely the most ecologically and economically important form of diazotrophic symbiosis occurs in specialized root structures called nodules, exclusive to the “nitrogen-fixing clade” of flowering plants (Soltis et al., 1995; Werner et al., 2014; Kates et al., 2022). Leguminous plants (Fabaceae) are the most familiar, with more than 90% of species able to nodulate (Sprent, 2007), but more sporadic nodular symbiosis occurs among nine further families (Kates et al., 2022).

Questions remain regarding the details of plant-bacterial partner matchup (Younginger and Friesen, 2019), but at the broadest scale partner choice is characteristic for higher level clades both of plants and of bacteria. Along with the genus *Parasponia* in Cannabaceae, the legumes form associations with α- and β-Proteobacteria (e.g., the genus *Rhizobium*, its segregates [e.g., *Bradyrhizobium, Sinorhizobium, Mesorhizobium*], *Burkholderia* and *Cupriavidus*); plants bearing this form of the symbiosis are termed “rhizobial.” Most prevalent in these plants, particularly in the North Temperate region, is an association with Rhizobiaceae of α-Proteobacteria (Sprent, 2009; Bellés-Sancho et al., 2023). While still the subject of active research, associations with β-Proteobacteria have been reported in a minority of plant taxa, primarily mimosoids of dry tropical regions, particularly of the Americas (Moulin et al., 2001; Gyaneshwar et al., 2011; Bontemps et al., 2016; Bellés-Sancho et al., 2023), and to our knowledge have yet to be reported in the Southeastern USA. In addition to rhizobial plants, the remaining eight plant families are thought to partner exclusively with Actinobacteria of the genus *Frankia* (Santi et al., 2013), with four genera widespread in the Southeastern USA (*Alnus, Ceanothus, Elaeagnus,* and *Morella* [=*Myrica*]), and are collectively termed “actinorhizal” plants.

Previous investigations have asked the relatively straightforward question of how specific the association is between nodulating plants and their diazotrophic symbionts, and the closely related question of how this association results in co-evolutionary pressures between the two partners. But answering this basic pair of questions has proven to not be straightforward (Sprent, 1994; Simms and Taylor, 2002; Martínez-Romero, 2009). Experimental systems and modeling studies have established partner choice as central to nitrogen-fixing symbiosis, ultimately rooted in selection against unbeneficial symbionts (Simms and Taylor, 2002; Heath and Tiffin, 2007, 2009; Batstone et al., 2020, 2022), but empirical investigations set in natural systems have been more equivocal (Parker, 2015). Part of the issue is that different studies can vary in their definitions of, and methods for quantifying co-evolution (Blasco-Costa et al., 2021). Equally important is that experimental and natural systems studies can both differ in spatial, environmental, and phylogenetic scale, which likely impacts the answer to these questions (Graham et al., 2018). How specificity is measured is particularly important, as the relevance of host choice seems obviously related to the degree of evolutionary divergence among strains. It may be that much of the difference among studies may boil down to how one conceptualizes measuring specificity, especially in a phylogenetic context. That is, specificity may be better understood as a phylogenetically structured property that emerges only beyond a certain evolutionary divergence, rather than as strict species-level exclusivity. This may suggest that there is an inflection point beyond which taxonomic specificity is relevant to consider. Therefore, it is imperative that research is structured to contain both spatial and phylogenetic components be leveraged (e.g., Pantinople et al., 2023) in order to investigate symbiont specificity.

The general lack of direct consideration of paired plant-microbe associations (e.g., Tamme et al., 2021; Doby et al., 2022; Sepp et al. 2023) is a key shortcoming of prior work for the specific purpose of understanding symbiosis. Such paired associations can help provide a basis for potentially better understanding and decoupling the phylogenetic and ecological factors, and ultimately the processes, at the core of the symbiosis. In sum, research on whether diazotrophs share the same distributional patterns with their host plants, particularly across contrasting ecosystems, remains essentially unknown. This gap means it is impossible to distinguish the factors that individually determine nodulating plant and diazotroph distributions from those that relate to the symbiosis itself, independently of, or interactively with, individual host/symbiont factors. A key next step is to distinguish among organism-specific assembly factors, and those pertaining to symbiosis is to assemble both at once and ask which better explains the current distribution of nodulating symbiosis. In addition to environmental and host drivers, biotic interactions that may potentially shape patterns of diazotroph community assembly remain to be poorly characterized. Simultaneous characterization of associated fungal communities has largely been unimplemented, and very little is known about the diversity and composition of fungal endophytes in root nodules. Uncovering the relationship between fungal communities and nitrogen-fixing bacteria will provide insights into how microbial assemblages are structured within the nodule and ultimately allow us to better understand the complex multi-kingdom dynamics that may influence biological nitrogen fixation.

High-throughput sequencing data from root-nodule symbioses, techniques best capable of producing a large-scale dataset for answering the questions we pose, have mostly been unutilized or excluded from large-scale investigations of diazotrophic composition, diversity, and distribution, since previous large geographic surveys have primarily used culture-based techniques (Doignon-Bourcier et al., 1999; De Meyer et al., 2011, 2015; Aserse et al., 2012, 2013; Bontemps et al., 2016). Here, we leverage a spatially and phylogenetically broad sampling from the local soil community to the immediate rhizosphere, root, and finally nodules themselves, to make progress in disentangling overall plant and microbial diversity from those aspects uniquely associated with symbiosis. We focus on southeastern North America, a rhizobial diversity hotspot (Sepp et al., 2023) with diverse communities of host nodulating plants. We hypothesize that symbiotic diazotroph diversity and composition are jointly driven by (1) habitat factors, which reflect available species to engage in symbiosis, and (2) plant host identity and phylogeny, which reflect host-symbiont alignment. Support for the first two hypotheses would suggest that nodulating plants possess adaptations specific to local diazotroph communities, and therefore we would expect that geographic origin (native vs nonnative) should also affect symbiotic diazotroph composition. We would predict therefore that (3) indigenous nodulating plants of the Southeast should recruit significantly different symbionts from co-occurring exotic nodulators. We also hypothesize that (4) native and nonnative host plants will exhibit variable bacterial-fungal co-occurrence patterns.

## MATERIALS AND METHODS

### Sampling scope

We focused on obtaining as many nodulating species as possible within field sites in an approximate transect of the southeastern United States. At 68 sites in Mississippi, Alabama, and north Florida, occurring approximately in a northwest to southeast transect (Figure 1), we sampled locally occurring nodulating species as possible, with 96 species sampled in this survey (see Appendix S1). Overall, we focused our sampling to ensure representation of the major lineages of nodulators that occur in the southeastern US, including all naturally occurring actinorhizal genera in the study area (*Alnus*, *Ceanothus, Elaeagnus*, and *Morella*) and most legume tribes (*n* = 18, from ca. 36). Sites were selected to maximize both species coverage, and major habitat types for nodulating plants, such as pineflats, prairie, and riparian environments. Sites were also selected to represent both intact native habitat and disturbed areas containing nonnative species. Mississippi and north Florida were the particular focus of habitat representation, with representation of mesic and dry habitats, and open and closed canopies. For several widespread species (*Centrosema virginiana*, *Chamaecrista nictitans*, *Crotalaria spectabilis*, *Desmodium rotundifolia*, *Galactia regularis*), 2-6 separate sites were sampled as an initial investigation of population variation in nodule symbionts, with 129 accessions overall. Few non-nodulating legume species occur in the study area; as a comparison, one non-nodulating species (*Senna obtusifolia*) was sampled.

**Figure 1.**
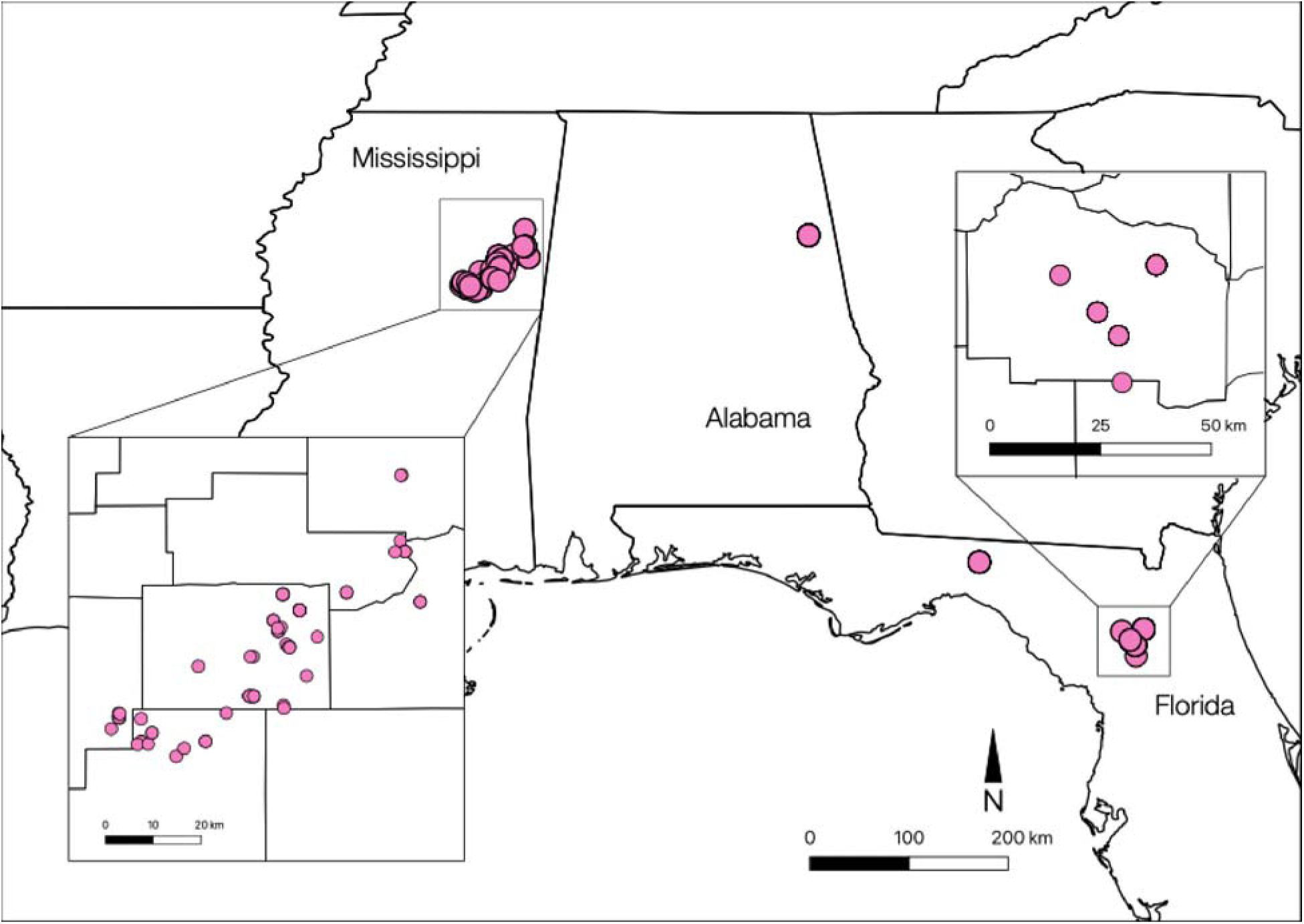
Map of 68 sampling locations in Mississippi, Alabama, and Florida, USA. Inset shows a finer scale map of sites within Mississippi and Florida where denser samples were collected for this study. Map generated using the QGIS Software (v3.24; QGIS Development Team 2021).

### Field sampling protocol

Plants were removed in entirety in an approximately foot-square clod with shovels that were washed between uses; the plant connection was maintained to ensure correct identification. These materials were transferred into the lab for sorting into four sample levels. First, soil was taken from clods at a distance ∼10-15 cm from the host plant; to reduce field contamination, soil clods were broken to take the interior of the sample. These samples of total soil near the host plant are referred to as *soil samples* hereafter; at sites with multiple species sampled, multiple soil samples were correspondingly taken. Roots and associated material adhering to roots were then dissected out of soil with clean forceps and material with verified nodules was transferred to microcentrifuge tubes. Nodules were confirmed by morphology in comparison with the literature, particularly field characteristics given (Sprent, 2009); the more unusual nodule morphology in actinorhizal species was particularly compared against photographic evidence in published literature. Plants were taken at peak green and preferably flowering as other seasons increase the risk of sampling senescent nodule organs. Prior to lab work, most samples were stored in absolute ethanol; for a subset of samples we investigated the use of a plant drier to desiccate samples (see Johnson, 2019; Pantinople et al., 2024; and Results).

### DNA extraction

Under a sterilized PCR hood, molecular-grade water was added to root samples, which were vigorously tapped to remove visible soil particles. The eluate from this step, which comprises the immediate environment of the root, was spun down for 5 minutes at 10,000 g; the pellet was reserved as the *rhizosphere sample.* The remaining root material was subjected to successive one-minute washes in 70% ethanol and 5% bleach to remove epiphytic organisms and environmental DNA based on Johnson (2019), washed twice with molecular-grade water to remove residual bleach, and then nodule material was dissected out from root material on a clean petri dish. Root material without nodules will be referred to hereafter as *root samples*, and dissected nodules will be referred to hereafter as *nodule samples.* In total, four sampling levels were investigated per collection event: *soil, rhizosphere, root,* and *nodule*.

A column-based open-source protocol (Williamson et al., 2014) to extract DNAs from tissue was optimized for nodule tissue extractions. Briefly, the dissected tissue and soil samples were transferred to a 2 mL tube containing 500 μL of lysis buffer (see Williamson et al. 2014) and processed with 4-6 sterilized metal beads in a tissue disruptor (Fisher Brand Bead Mill 24) using the following program: speed = 3.10 m/s, number of cycles = 04, cycle time = 30 s, pause = 5 s to lyse the cells and release the cell contents in the buffer solution. The tubes were incubated for 20 minutes in a hot water bath at 65°C and processed in the tissue disruptor a second time to ensure complete lysis of cells. The tubes were then incubated for 10 minutes in the hot water bath at 65°C and then centrifuged at 10,000 g for 2 minutes. The supernatant was transferred to another set of tubes containing 200 μL of potassium acetate (49.08%) to precipitate proteins and detergents. The tubes were stored in a −20°C freezer for at least an hour before proceeding to purify the dissolved DNA. For purification, tubes with precipitated impurities were centrifuged at 10,000 g for 30 minutes, and the supernatant was transferred and mixed with 1.2 mL guanidine hydrochloride (31.5 g guanidine hydrochloride in 316.5 mL ethanol and 152 mL molecular grade water). The mixture was transferred to a silica spin column (5 μl EconoSpin® columns, Epoch Life Science, Missouri City, Texas, USA), 700 μL at a time, and centrifuged at 10,000 g for 2 minutes, discarding the flow through. The column was washed twice with 500 μL wash solution (10 mL 1M Tris at pH 8.0, 2 mL 0.5 M EDTA, 10 mL 0.5 M NaCl, 670 mL ethanol, and 308 mL molecular grade water) then once with 80% ethanol, centrifuging at 10,000 g for 2 minutes for each wash. The column was centrifuged at 10,000 g for 5 minutes to dry and the DNA was eluted off of the column using 200 μL of elution solution (5 mL 1 M Tris at pH 8.0 in 495 mL molecular grade water) after a 10-minute incubation at room temperature, and finally centrifuged at 10,000 g for 2 minutes. Tubes containing the eluate were quantified via Qubit (ThermoFisher Scientific, Waltham, Massachusetts, USA) prior to PCR. This protocol has the potential to be broadly useful because it avoids the need for automated extraction robots, 96-channel pipettors, and other specialized expensive equipment as called for in the most recent comparable protocol (Johnson, 2019) but available in few labs; see the full lab protocol in Appendix S7.

### PCR amplification

For bacterial characterization, we amplified the V4 region of 16S ribosomal DNA using the primers 515f and 806r (Thompson et al., 2017). We applied the following thermal cycling conditions: initial denaturation at 95°C for 3 mins, followed by 35 cycles of 95°C for 45 s, annealing at 52°C for 1 min, and 72°C for 1.5 mins, and a final elongation at 72°C for 10 mins. We then normalized the concentration of successful PCR products before submitting to the Michigan State RSTF core for 250 bp paired-end reads sequencing on an Illumina MiSeq using a one-step amplification protocol (Kozich et al., 2013). Except where stated, all amplification procedures were carried out using DreamTAQ Mastermix (ThermoFisher, Waltham, Massachusetts, USA) and primer concentrations of 0.5 μM. Filter pipette tips were used, and to reduce contamination, each PCR hood was bleach- and UV-sterilized before each use.

For fungi, we amplified the fungal ITS1 region using the primers ITS1-F (Gardes and Bruns, 1993; chosen to reduce off-target host DNA) and ITS2 (White et al., 1990). We first verified the presence of amplifiable DNA by directly using the primers and the following thermal cycling conditions: initial denaturation at 95°C for 3 mins, followed by 35 cycles of 95°C for 45s, annealing at 50°C for 1 min, and 72°C for 1 min, with a final elongation at 72°C for 10 mins. Then, we re-amplified successful products using the primers tagged with 5’ end overhangs with the following thermocycler protocol: initial denaturation at 95°C for 5 mins, followed by 30 cycles of 95°C for 30 s, annealing at 52°C for 30 s, and 72°C for 30 s, with a final elongation at 72°C for 5 mins and primer concentrations of 0.1 μM. Initial PCR optimizations used 30 cycles and annealing temperatures of 55°C; this was later optimized to 35 cycles and annealing temperatures of 50°C; coding this as a factor we found that these parameters increased success but did not affect standard diversity metrics. Amplicons were also submitted for sequencing, and wet lab protocols to avoid contamination were done similarly as those for bacteria.

### Sequence processing and analysis

To identify taxa, amplicon data were analyzed within the QIIME 2 package version 2024.2 (Caporaso et al., 2010; Estaki et al., 2020) following recommended parameters using the most up-to-date Greengenes (McDonald et al., 2012) and UNITE (Nilsson et al., 2019) reference databases for bacterial 16S reads and fungal ITS reads, respectively. A trial of Greengenes2 (McDonald et al., 2023) was also made; however, the decision to remove eukaryotic 18S in this version prevented our ability to identify host DNA and therefore its use was infeasible (see below). The sequence reads were denoised using Dada2 (Callahan et al., 2016) to correct for errors, merge paired-end reads, and remove PCR chimeras. Primers were also removed in the process by trimming the primer length from the 5’ end, and on the 3’ end of the R2 reads based on FastQC (Andrews, 2015) quality plots. Clustering of the Greengenes and UNITE databases were set at 97% and 99% identity threshold, respectively. We then ran the merged reads against Greengenes and UNITE using the “sklearn” module (Estaki et al., 2020) in QIIME 2 for taxonomic classifications. To efficiently filter out host plant DNA from each of the genetic loci, we applied optimization strategies from Pantinople et al. (2024) for endophytic tissues. Briefly, we removed host chloroplast and mitochondrial 16S by taxonomic filtering within QIIME2 using annotated chloroplast and mitochondrial operational taxonomic unit (out) classifications (*--p-exclude c Chloroplast, f mitochondria*), while we filtered out host ITS sequences using annotated genus level OTUs.

### Diversity and composition analysis

In order to characterize microbial diversity, we calculated standard species and phylogenetic diversity metrics within QIIME 2. We used Shannon entropy as a measure of overall species diversity and Faith’s phylogenetic diversity (PD) to quantify diversity based on the total length of phylogenetic branches connecting members of the community. Sequence rarefaction cutoffs were set for both diversity statistics based on generated rarefaction plots (see Appendix S2), taking into consideration the threshold that best represents microbiome diversity while still maintaining a sufficient number of samples for downstream analyses. Nodule, root, and rhizosphere samples showed saturation in rarefaction plots at 100 reads; this makes sense since these are low-diversity samples and nodules in particular tend to be infected by a single strain of symbiont at >90% frequency. By contrast, soil samples are much more diverse (∼20X OTU count) and do not exhibit saturation until 200 reads. Thus, these cutoffs (100 and 200 reads) were implemented for downstream analyses depending on sample type. Samples were usually successful, but rhizosphere samples, which were strictly defined as soil material adhering to the root and nodule samples, were more likely to be excluded, consistent with the low biomass and often sub-5 ng yield of these samples.

To identify variation in microbial community composition, we generated Jaccard distance matrices within QIIME 2 and performed PERMANOVA using the *beta-group-significance* function in the same platform to test for statistical significance. We also performed differential abundance analyses using the Analysis of Composition of Microbiomes (ANCOM; Mandal et al., 2015) function implemented within the QIIME 2 platform. ANCOM is a statistical method that helps identify taxa that significantly vary in relative abundance across groups, often visualized in a volcano plot. The x-axis represents the effect size (clr; centered log-ratio) while the y-axis represents statistical significance (W statistic). Points in the top right corner of the plot represent taxa that are most differentially abundant and highly significant. We ran Mantel correlation tests between nodule UniFrac distances and host patristic distances using the “vegan” package (v.2.6.4; Oksanen et al., 2022) in R, to see if microbiome UniFrac distance is associated with host phylogenetic distance. The host patristic distances were extracted from a host plant phylogeny we constructed using the package “rtrees” (Li, 2023) in R.

To evaluate and visualize host-symbiont associations, we performed bipartite network analyses using feature tables generated from QIIME2 as input data. We selected the top 20 symbiont genera based on total abundance across samples. Mean relative abundance were then calculated for each host-symbiont pair and the pairs with ≥ 0.008 abundance were retained for network construction. Bipartite networks were generated in python using the NetworkX package (Hagberg et. al., 2008) and visualized using matplotlib (Hunter, 2007) where host and symbiont were defined as two distinct node sets. Node size was scaled by relative abundance and edge width represented association strength.

### Co-occurrence network analysis

To characterize bacterial-fungal associations in the nodule, we performed a cross kingdom co-occurrence network analysis with merged 16S and ITS feature tables (with taxonomy) from QIIME2 as input data. For the network, we selected the top 30 bacterial and fungal genera based on mean relative abundance and only included rhizobial symbionts. Microbial associations were estimated using Spearman’s rank correlation coefficient (Wisniewski and Brannan, 2025) and only significant associations with an absolute correlation coefficient ≥ 0.25 were kept for network construction. Networks were generated in python using the NetworkX package (Hagberg et. al., 2008) and visualized using matplotlib (Hunter, 2007), with node size scaled by mean relative abundance and edge width and color representing correlation strength and direction, respectively.

### Diazotrophic determinations

Here we followed a strategy used by several previous studies of diazotrophs (e.g., Sepp et al., 2023; Li et al., 2025) using taxonomic determinations to infer function, in this case diazotrophic competency. Diazotrophic determinations followed literature sources including Nichio et al., 2025 and particularly the ICSP taxonomic database (International Committee for Systematics of Prokaryotes; https://sites.google.com/view/taxonomyagrorhizo/). Because this approach infers diazotrophic function rather than measuring it, it is possible diazotrophic strains could be overcounted because diazotrophic variation is known below the genus level. However, this concern is partly mitigated by the use of multiple sampling levels (i.e., soil, rhizosphere, root, and nodule). A near mono-infection (e.g., abundance >90%) of a nodule by a single diazotrophic genus, seen in most samples, was regarded as likely evidence that the nodulating bacterium was viewed as evidence of the probable diazotroph. For the exception, concerning actinorhizal taxa, see Discussion.

## RESULTS

### Relative abundance and composition

After sequence denoising, we recovered a total of 8,555,117 high-quality 16S sequencing reads from 360 samples across all sampling levels (mean 23,764 reads per sample) sorted into 38,283 total bacterial OTUs (with a 97% sequence identity cutoff). For fungi, we recovered a total of 3,813,457 high-quality ITS sequencing reads from 249 samples across all sampling levels (mean 15,315 reads per sample) grouped into 2,523 total fungal OTUs (with a 99% sequence identity cutoff). From just the nodule samples, we recovered a total of 6,257,267 16S reads (mean 41,438 reads per sample) from 151 samples with 3,731 total bacterial OTUs, and a total of 1,767,550 ITS reads (mean 13,808 reads per sample) from 128 samples with 1,062 total fungal OTUs. We tested for an effect of preservation method (ethanol vs. desiccation) and sequencing batch on Faith’s PD and Shannon diversity, finding no methodological effect. Our sequencing results confirm the presence of both bacterial and fungal communities that coexist within host plant nodules and demonstrate that bacteria are dominant symbionts co-occurring with a relatively less diverse fungal microbiome.

The most dominant bacterial symbionts belonged to Alphaproteobacteria, which are found in highest relative abundance across all sampling levels (16 to 89%). Other bacterial classes including Gammaproteobacteria, Deltaproteobacteria, Betaproteobacteria, and Actinobacteria follow, each constituting less than 20% of relative bacterial abundance among samples (Figure 2A). At a finer taxonomic level, **we recognize nine bacterial genera** as potential diazotrophs based on findings from primary literature (see Methods). The most abundant was *Bradyrhizobium*, found across all sampling levels and following a trend of increasing relative abundance from surrounding soil to nodule (1 to 49%; Figure 2B). The same pattern was observed in *Rhizobium*, *Mesorhizobium*, and *Agrobacterium* where most of the abundance were found in roots and nodules, and together account for more than 30% of the nodular microbiome. *Azorhizobium* was mostly found in the rhizosphere (2.8%) and nodule (2.4%), *Burkholderia* was most abundant in the rhizosphere (6.3%), and *Sphingomonas* was mostly in the roots (2.4%). *Sinorhizobium* and *Frankia* were found virtually only in roots and nodules, respectively. Overall, diazotroph relative abundance showed an increasing trend as sampling level approached the nodule organ.

**Figure 2.**
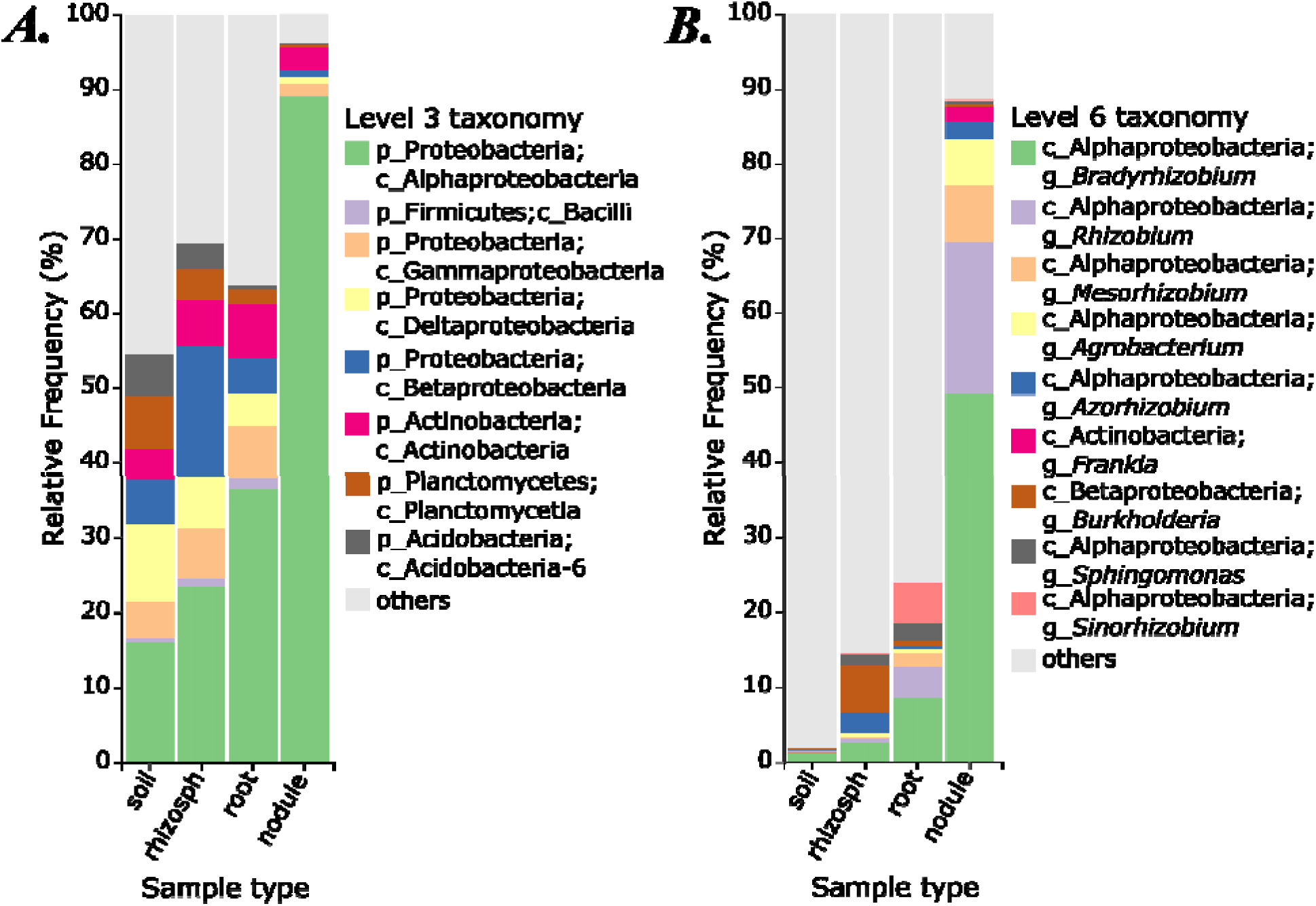
Relative frequencies (%) of (A) dominant bacterial classes and (B) major diazotrophic genera in colors, filtered out of the broader community (gray) across the four sampling levels (soil to nodule; rhizosph = rhizosphere). Data from aggregated 360 samples and 84 plant species from MS, AL, and FL (USA).

For fungi, Agaricomycetes dominated the rhizosphere (43%), root (34%), and nodule (32%), while Leotiomycetes was primarily abundant in soil (47%; Figure 3A). Sordariomycetes, Dothideomycetes, Eurotiomycetes, Tremellomycetes, and Saccharomycetes were relatively less abundant (<30%) but were found across all sampling levels. At the genus level, *Russula* dominated the nodule (10%; Figure 3B), while *Microglossum* was the primary fungus in soil (44%) and was undetected in other sampling levels. *Fusarium*, *Cladosporium*, and *Curvularia* were found relatively less abundant (3 to 9%) across samples but were virtually absent in soil.

**Figure 3.**
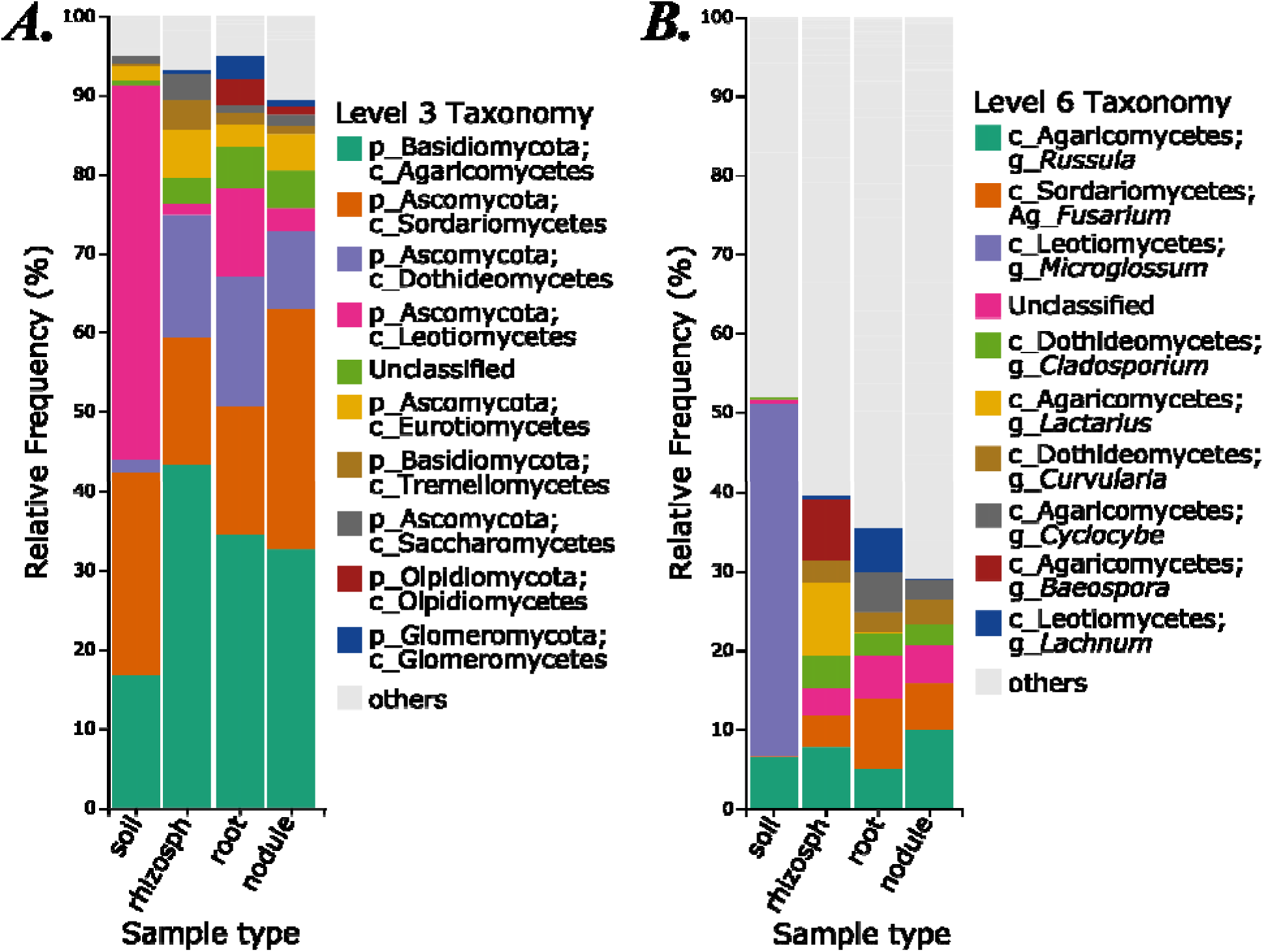
Relative frequencies (%) of (A) dominant fungal classes and (B) genera filtered out of the broader community (gray) across multiple sampling levels (soil to nodule; rhizosph = rhizosphere). Data from aggregated 249 samples and 84 plant species from MS, AL, and FL (USA).

Our results demonstrate that plant nodules host a complex set of microbial communities including putatively non-diazotrophic bacteria and fungi that coexist and interact with nitrogen fixers. Warranting further investigation is the potential unknown role of co-occurring symbionts in biological nitrogen fixation and other nodular processes.

### Diversity and composition of nodule microbiome across habitat

Host plants found in prairies had by far the most diverse nodular communities across habitat types (Figures 4A&B). Both Faith’s PD and Shannon diversity were significantly higher in prairies than ditches, roadsides, and wood edges (pairwise Kruskal-Wallis, *p* < 0.05 for all three). Prairies, bottomland woods, flat woods, pine woods, and calcareous prairies had the highest symbiont diversity, while roadsides, fields, upland woods, wood edges, and ditches had the lowest (Figures 4A&B), revealing a general pattern of lower diversity in disturbed versus natural habitats. However, across all habitats, both nodular Faith’s PD and Shannon diversity were not statistically different (Kruskal-Wallis, *p* = 0.17 for both diversity metrics).

**Figure 4.**
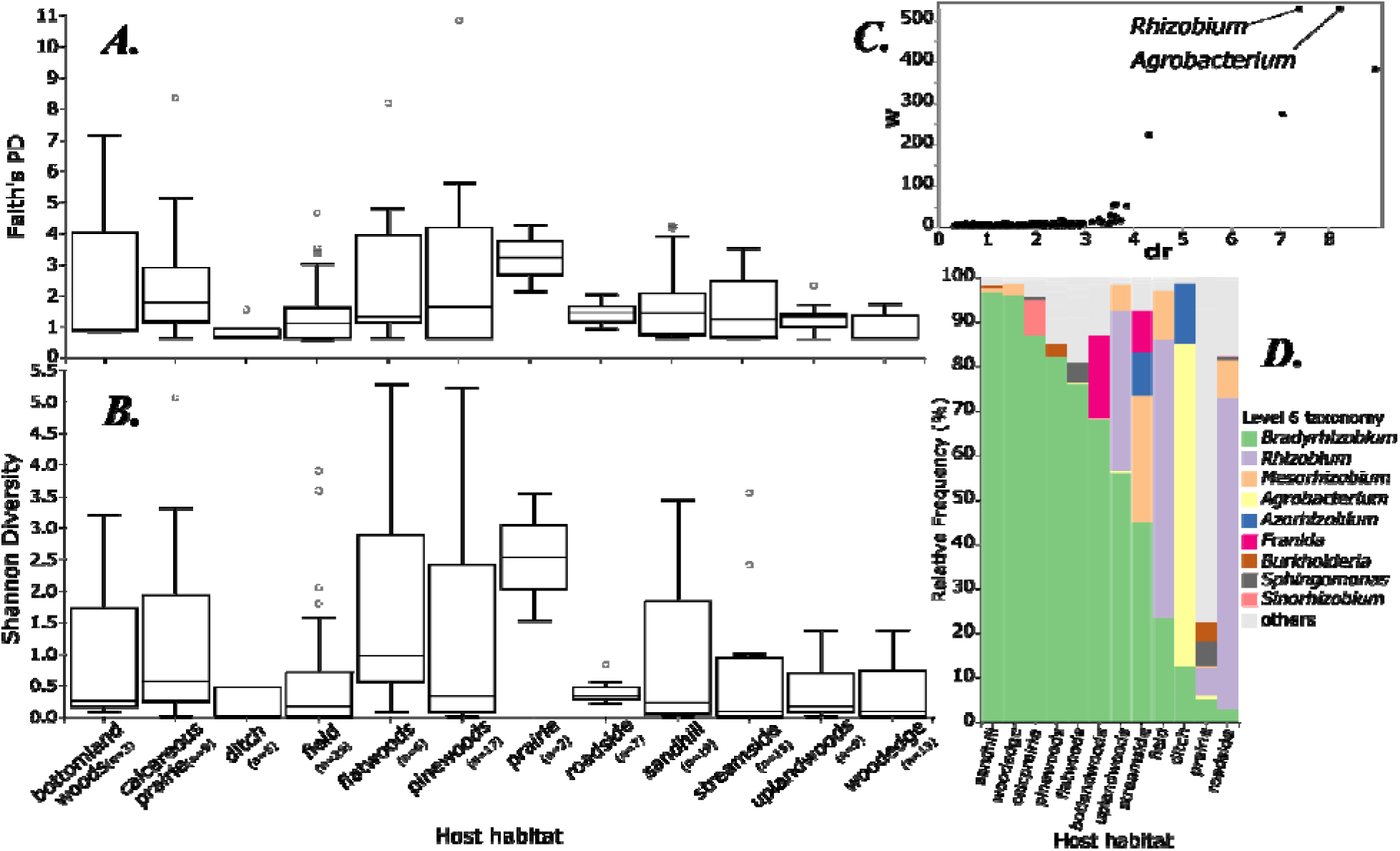
Bacterial diversity and community structure of nodulating host plants across habitat types. Boxplot of (A) Faith’s PD and (B) Shannon diversity of nodular bacterial communities. (C) ANCOM volcano plot of communities showing *Agrobacterium* and *Rhizobium* to be differentially abundant across host habitat (labeled points are significant). (D) Relative frequencies of major nodular diazotrophic genera. Reflecting the ANCOM result, disturbed field and roadside habitats were closely associated with *Rhizobium*, while wetter disturbed ditch sites were associated with *Agrobacterium*.

Our analyses based on Jaccard distances, however, revealed highly variable nodular community structures across habitats (PERMANOVA, *p* = 0.001). In addition, the abundance of both *Agrobacterium* and *Rhizobium* were detected to be significantly different in nodular communities across host habitat (Figure 4C), based on our ANCOM results; this reflects the association of *Agrobacterium* with the host *Sesbania* found in **semi aquatic habitats**, and the prevalence of *Rhizobium* in **ruderal habitats** (Figure 4D). *Agrobacterium* was more abundant in the nodules of hosts found in ditches while *Rhizobium* was more abundant in fields, prairies, roadsides, and upland woods (Figure 4D).

### Phylogenetic patterns of diversity and composition

Comparing across host species, we found no significant difference in both the Faith’s PD and Shannon diversity of their bacterial nodular communities (*p* = 0.25, 0.16, respectively; Appendix S3). However, looking at broader host taxonomic levels, our results reveal significant variation in both the standard diversity metrics across genera (*p*= 0.02, 0.03; Appendix S4) and tribes (*p* = 0.01, 0.02; Figures 5A & B). Thus, microbiome diversity differs among higher-level host clades, although not at the species level.

**Figure 5.**
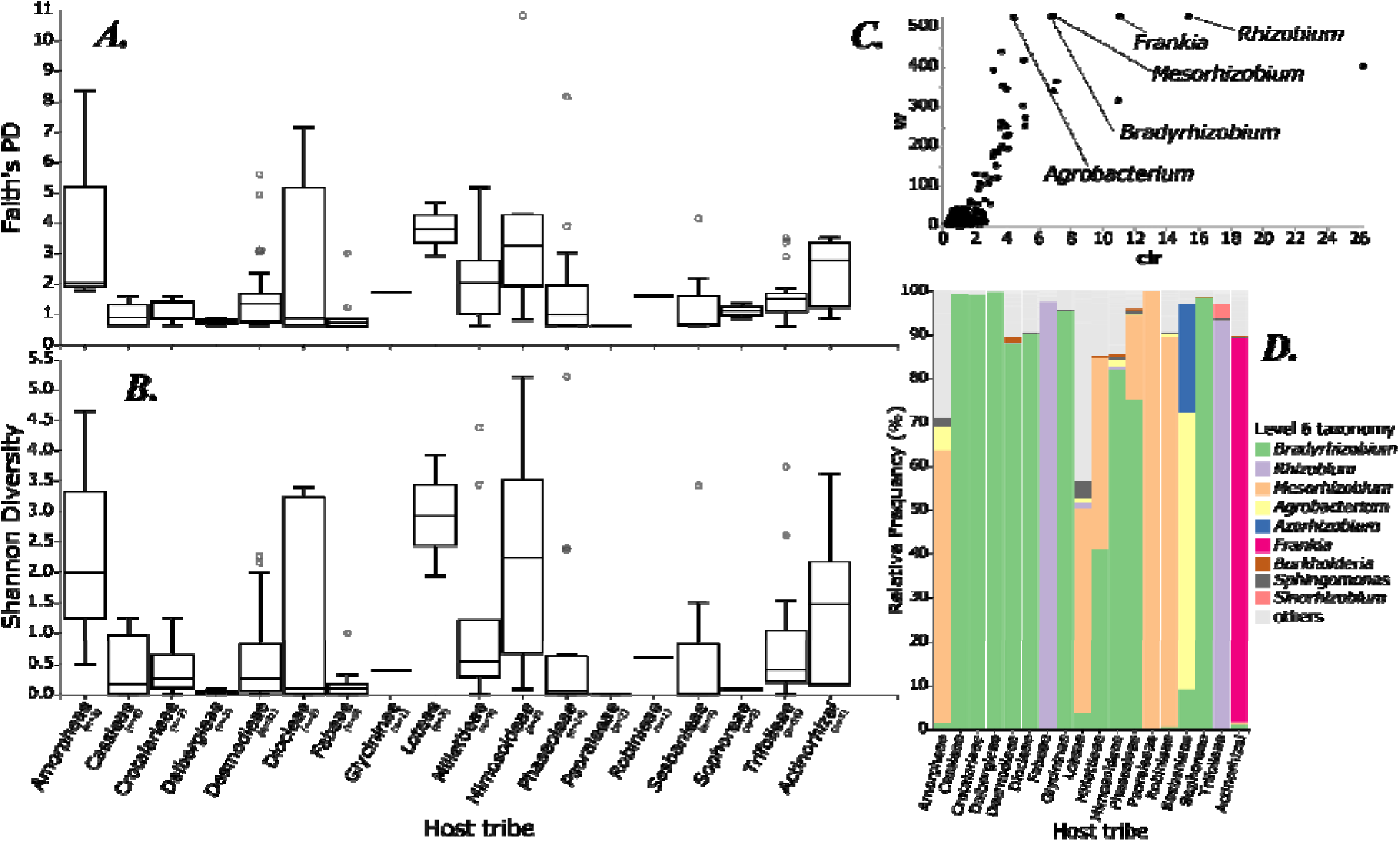
Bacterial diversity and community structure across nodulating plant host tribes. Boxplot of (A) Faith’s PD and (B) Shannon Diversity of nodular bacterial communities. (C) ANCOM Volcano plot of communities (labeled points are significant) showing *Bradyrhizobium*, *Sinorhizobium*, *Frankia*, *Rhizobium*, and *Mesorhizobium* to be differentially abundant across host tribes. (D) Relative frequencies of major nodular diazotrophic genera across host tribes.

Because few species of actinorhizal plants occur in the study area, these are treated as a unit. Actinorhizal plants and most legume tribes are primarily characterized by a single dominant diazotrophic genus. Primarily native tribes are associated primarily with *Bradyrhizobium* and *Mesorhizobium*; primarily non-native tribes (Fabeae, Trifolieae) are associated with *Rhizobium*.

In terms of differential abundance, there are highly variable community structures among host tribes as indicated by our ANCOM results (Figure 5C). The abundance of *Frankia*, *Bradyrhizobium*, *Rhizobium*, *Mesorhizobium*, and *Agrobacterium* were detected to be significantly different across host tribes (Figure 5D). We also found a significant correlation (*p* = 0.015) between host phylogenetic distance and nodular bacterial community distance, suggesting that closely related hosts tend to have a similar set of bacterial microbiomes in the nodules (Appendix S6). Our host-symbiont bipartite network further revealed the dominance of *Bradyrhizobium* and its association across multiple host tribes, indicating broad host ranges (Figure 6). However, majority of diazotrophic genera including *Rhizobium*, *Agrobacterium*, *Sinorhizobium*, and *Azorhizobium* only strongly associate with only one to a few sets of host tribes which further supports our previous finding suggesting a level of host specificity among these symbionts.

**Figure 6.**
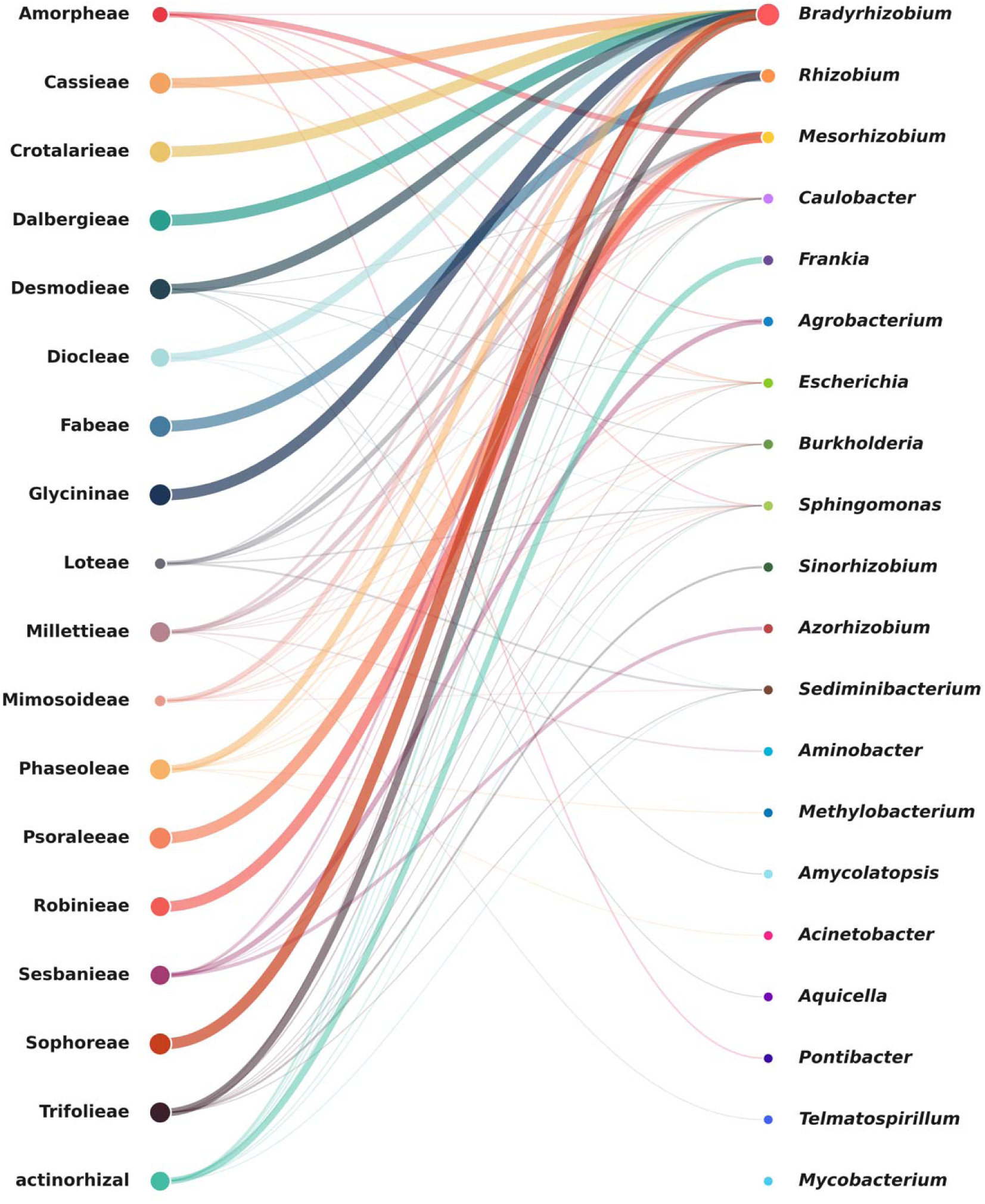
Bipartite network of host plant vs. bacterial symbionts. Nodes on the left represent host tribes and nodes on the right represent the top 20 bacterial genera. Node size and edge (link) width are scaled by mean relative abundance. Links are filtered at a 0.8% (0.008) threshold to highlight significant symbiotic associations. Edges are colored by the source host tribe.

### Microbiome diversity and composition between native and nonnative hosts

Our results show that bacterial Faith’s PD and Shannon diversity did not vary significantly across indigenous and exotic host plant nodules (Fig. 7A&B), suggesting that the plants harbor similarly diverse nodular bacteria regardless of distributional range. However, in terms of community composition, our analyses based on Jaccard distances revealed significant differences within nodules and roots (PERMANOVA, *p =* 0.001 for both), but not in rhizospheres and soils (PERMANOVA, *p =* 0.325 and *p =* 0.549, respectively). The taxa barplot (Fig. 7D) also showed contrasting community structure revealing *Rhizobium* to be a highly dominant diazotroph in nonnative host plant nodules compared to the native group. The ANCOM result further supported this finding, demonstrating that *Rhizobium* is significantly differentially abundant across the two groups of nodulating host plants (Fig. 7C), specifically enriched in nonnatives. This is an interesting result because we did not see *Rhizobium* specifically enriched in neighboring native plants and their local rhizosphere. We found no significant difference in the ANCOM abundance in soil, rhizosphere, and root between native and nonnative host plants (Appendix S5). Our Jaccard and ANCOM results suggest that nonnative host plant nodules may have a closer association with *Rhizobium* and specifically recruit this diazotroph in their nonnative range, regardless of the available surrounding microbial species pool.

**Figure 7.**
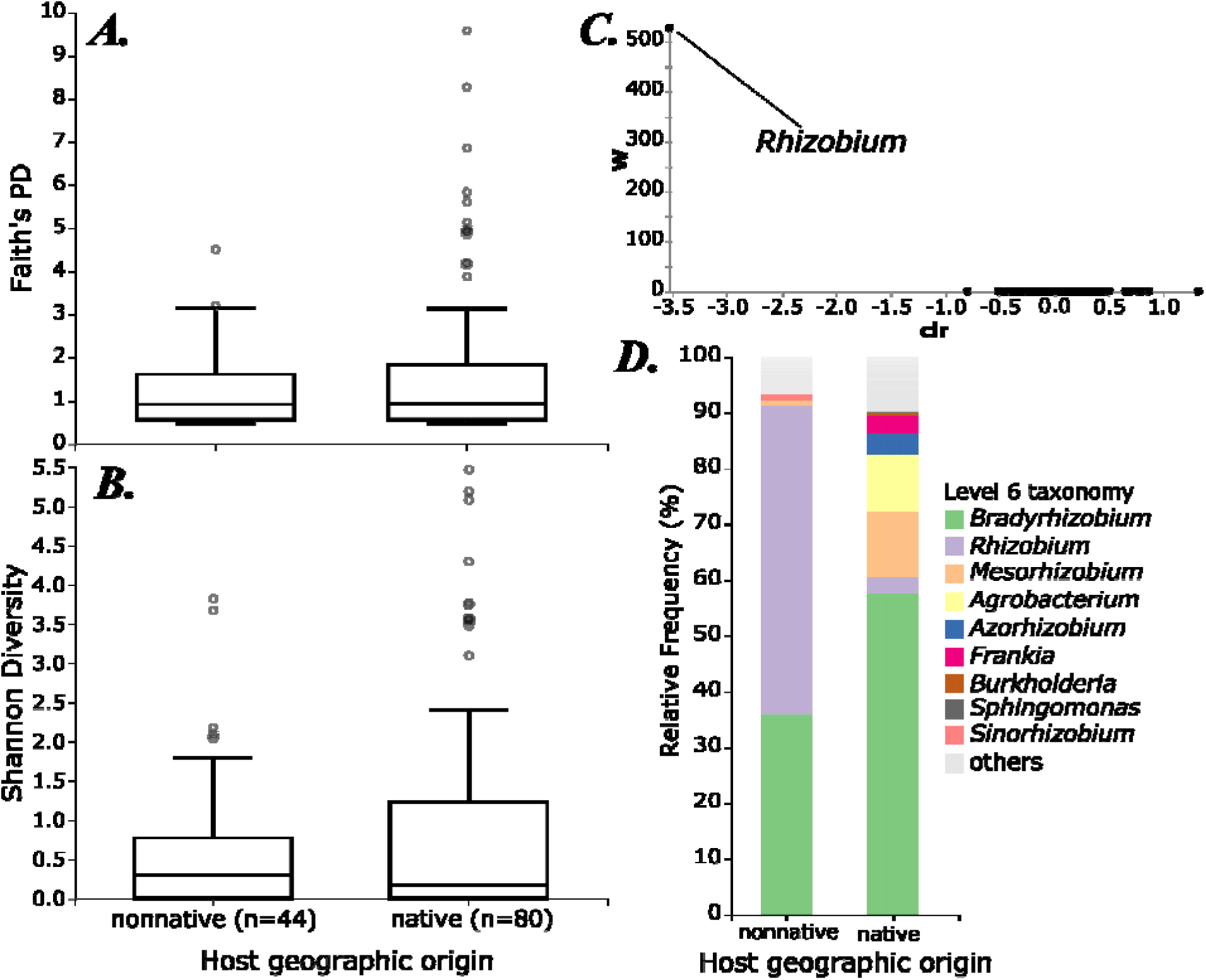
Bacterial diversity and community structure between native and nonnative host plant nodules. Boxplot of (A) Faith’s PD and (B) Shannon Diversity of nodular bacterial communities, showing no significant differences. (C) ANCOM volcano plot of communities across host geographic origin, labeling only significant points. This analysis shows *Rhizobium* to be the sole differentially abundant taxon, being closely associated with the majority of nonnatives in the study (see also Fig. 7d). (D) Relative frequencies of major diazotrophic genera between native and nonnative hosts, showing *Rhizobium* being particularly enriched in nonnatives.

### Co-occurrence patterns between native and nonnative hosts

Nodular bacteria-fungi co-occurrence networks differed between native and nonnative host plants. The native host plant network comprised a total of 161 significant associations, whereas the nonnative host plant network contained 283 associations. Positive correlations accounted for the majority of all associations, with 95.6% and 90.1% in native and nonnative hosts, respectively. Compared to the native host network, the nonnative network had a greater number of negative correlations across and within microbial groups.

## DISCUSSION

### Fuzzy specificity predominates in the Southeastern legume flora

Through application of ANCOM at successive collapsing levels, we found that our measure of specificity does not occur at low phylogenetic levels (here at the level of species) but is significant for clades above the species. This result was specific to diazotrophic genera (Figure 5C). Similarly, a Mantel test of host patristic distance vs. community composition was significant, suggesting this was not simply a statistical power effect. Our results were consistent with the concept of “fuzzy” specificity; that is, related nodulating plants host similar sets of bacteria that are related primarily at the clade level rather than the species or strain level. This concept, also often described as “phylogenetic host specificity,” has been described in parasites from viruses to arthropods (Cooper et al., 2012; Santiago-Alarcon et al., 2014; Park et al., 2018; Gupta et al., 2020), in mutualisms such as mycorrhizae (Tedersoo et al., 2013; Dong et al., 2021), and perhaps most clearly in the coral-Symbiodiniaceae mutualism (Rouzé et al., 2019) where specificity signal seems specifically restricted to higher level clades.

In terms of mechanism, fuzziness should arise (1) from a co-occurring duet of co-speciation and host-switching (Rouzé et al., 2019) which could result in relatively symmetrical fuzziness among host and symbiont. However, (2) conservatism of functional traits relevant to symbiosis could conceivably also contribute (Yang et al., 2012). This would lead to the prediction of more fuzziness in the less conservative partner, as well as (3) differing speciation-extinction regimes among host and symbiont (Cooper et al., 2012), which should create fuzziness for only the higher net-diversification partner. To our knowledge, this concept has yet to be directly studied in the nitrogen-fixing symbiosis but Parker (2015) discusses a limited degree of phylogenetic conservatism evident in the host associations of the diazotroph *Bradyrhizobium*.

The idea of fuzzy specificity – that plant-diazotroph specificity operates primarily at higher phylogenetic levels – is intuitive in the nitrogen-fixing symbiosis as an emergent property arising from key evolutionary trade-offs between generalist and specialist nodulators (Ehinger et al., 2014; Batstone et al., 2020). It would also be an intuitive explanation of differing results among studies focusing on whether nodulating symbiosis displays co-evolution (Martínez-Romero, 2009). The choice between these lifestyles is not straightforward given that the nodule organ is a costly plant investment in terms of photosynthate (Kaschuk et al., 2012). Microbial genome evolution is driven partly by nitrogen limitation (Shenhav and Zeevi, 2020) which could lead to nutritional conflict with the host. And not all plant-diazotroph pairs represent functionally equivalent symbioses (Batterman et al., 2018) so that plants and microbes are likely to find themselves in evolutionary conflict.

Correct match-up, associated with the highest efficiency (Sprent, 2009), is mediated by signaling factors such as *nodA*, which are essential for inducing infection (Bisseling and Geurts, 2020). Host-symbiont mismatch, by contrast, leads to poorly co-adapted plant-microbe pairs and inefficient net nitrogen fixation (Simms and Taylor, 2002; Friesen, 2012; Moyano et al., 2017). Indeed, the nodule organ itself has been interpreted as part of a strategy for plants to exert control over the symbiosis and eject underperforming symbionts (Chomicki et al., 2020). Nevertheless, plants are at pains to be flexible in partner choice when partners are limited, just as we would expect in disturbed or alien habitats (Parker et al., 2006; Simonsen et al., 2017). The foregoing considerations are compatible with both (1) limited but present host switching regimes (constrained perhaps by conservatism in methods of symbiont recognition) and (2) functional conservatism (constrained by the difficulty of acquiring a new nitrogen-fixing competence or symbiotic capability) as enumerated above. It is also likely that (3) diazotrophs differ from their hosts in speciation-extinction regime, although this is not necessarily obviously connected to nodulation.

Under these constraints, it would follow that specificity would occur within bounds partly delimited by evolutionary history (as a balance between flexibility among similar partners and discriminating choice when presented with unfamiliar partners), and perhaps more proximally the evolutionary history of gain of *nod* factors (one way in which host switching would be facilitated), an element that is feasible but remains to be investigated under this framework. An important next step would be to phylogenetically investigate *nif* and *nod* genes as representatives of the history of nitrogenase function and host recognition respectively, which would assist in differentiating (1) and (2).

### Nodulating plant clades differ in nodule diversity and composition

While we found no evidence for species-level consistent differences in nodule diversity, these differences were highly significant at higher phylogenetic levels, here measured at the levels of genus and tribe. Excluding outlier samples, the highest median phylogenetic diversity was observed in Amorpheae, Loteae, mimosoids, and actinorhizal plants (all > 3; Figure 5A).

Shannon diversity outliers were similar, with high values in the aforementioned four clades (all > 1; Figure 5B). This result is interesting as the nodule functions as a rarefied microbiome environment (Chomicki et al., 2020; de Faria et al., 2022); thus, high microbial diversity could be seen as negative in the sense of evincing a poorly controlled nodule environment. In this regard, the highest diversity nodule environments are notable as containing the phylogenetic outliers in the dataset. Most species we sampled are core papilionoid legumes, thus in the most diverse clade of nodulators. The actinorhizal category comprises all sampled plants that fall outside the legumes, the mimosoid group is the only non-papilionoid group sampled, and

Amorpheae and Loteae fall outside the core papilionoids (although we also sampled other divergent papilionoids that did not display high nodule diversity). Mimosoids are notable as already being documented as relatively promiscuous in host choice, including both alpha and beta rhizobia (Moulin et al., 2001; Bontemps et al., 2016) as well as relaxed infection modes (De Faria et al., 1988), and actinorhizal plants in general are thought to have less ability to impose host control over choice (Pawlowski and Sprent, 2008; Ardley and Sprent, 2021). Core papilionoids, by contrast, have more specific infection modes as well as direct antimicrobial activity against non-specific microbes (Eaker et al., 2025). Thus, our finding of highly significant phylogenetically structured diversity differences among tribes, points to host-level differences in microbiome control.

We also conducted ANCOM analyses to investigate phylogenetically structured host differences in microbial community composition. As mentioned above, ANCOM successfully recovered diazotroph differences at the host clade level. These were *Agrobacterium, Bradyrhizobium, Frankia, Mesorhizobium,* and *Rhizobium*, corresponding roughly to all of the diazotrophs with multiple samples and thus sufficiently powered to detect a difference. However, the analyses failed to discover any compositional differences in non-diazotrophic community members. This could indicate relatively casual nodule occupancy by unspecialized endophytes lacking in host specificity. This result differs from endophytes from other hosts and tissues, such as leaf endophytes, where bacterial endophyte composition has been repeatedly found to be significantly phylogenetically structured (Pantinople et al., 2024).

### Nodule diversity differs by habitat

Although overall differences were not found to be significant, nodular diversity was generally highest in habitats that also display high vascular plant diversity, namely bottomland woods, prairie, and sandhill habitats. The lowest nodule microbiome diversity was seen in the lowest-quality habitats, namely fields, ditches, and roadsides. The habitats in the study region are low-diversity overall and many predominantly contain non-native nodulating plants in the Fabeae and Trifolieae, thus also displaying restricted nodulator diversity.

Bacterial symbiont composition significantly varied across habitat. The ANCOM analysis recovered two taxa that were significantly enriched in certain habitats. *Rhizobium,* the first result, is likely confounded by the native/non-native contrast, as non-natives are enriched in disturbed habitats and the *Rhizobium* specificity is likely endogenous as we argue above. *Agrobacterium*, the other result, is probably an artifact of the GreenGenes backbone taxonomy we rely upon. The Rhizobiales (Hyphomicrobiales) have been the subject of recent taxonomic revision to resolve non-monophyly (Kuzmanović et al., 2022; Ma et al., 2023; diCenzo et al., 2024), and this has particularly affected the delimitation of *Rhizobium* vs. *Agrobacterium*, neither of which as traditionally delimited is monophyletic (Ma et al., 2023). Only isolates of *Sesbania herbacea* (associated with the wettest habitats in this study) were identified by the taxonomic classifier as belonging to this genus, likely making it a divergent isolate but in need of future further data as short amplicon fragments are not ideal for investigating novel taxonomic placements.

### Nodule diversity and composition differ by host geographic origin

We found evidence suggesting that host nodulators differentially recruit diazotrophs in the Southeastern US based on native status. Nodular bacterial composition differed significantly between native and nonnative host plants, with solely *Rhizobium* being at significantly higher frequency in exotic host nodules than in natives. This finding supports the idea that host plant nodules differentially recruit diazotrophs based on their original biogeographic context, especially given that surrounding bacterial community pools are similar in terms of composition for both native and nonnative hosts. More specifically, it seems most likely that this result reflects a binary in biogeographic histories comprising the modern Southeastern legume flora, which shapes differing optimal symbionts. Natural diazotroph diversity is not evenly distributed across Earth (Sepp et al., 2023). Native legumes in this area are nodulated by several clades but particularly *Bradyrhizobium (Sprent, 2009)*, which is prevalent in but not solely found in North American nodulators. Introduced legumes of eastern North America, however, are generally western Eurasian in origin, and most were brought over for specific purposes of forage and cover crop systems (selecting for weedy plants with short lifecycles) rather than originating as waifs (Norden and Kirkman, 2006). In this original range, most legumes are nodulated by *Rhizobium.* The foregoing facts are theoretically relevant to biogeography because species highly reliant on specialized symbionts should experience differing dispersal ecology as compared to flexible or non-symbiotic species. As has been argued previously (Parker et al., 2006; Simonsen et al., 2017), non-native and invasive plants cannot easily change symbiotic partners and therefore the range may be limited by the availability of appropriate symbionts in foreign soil environments. Experiments have shown that exotic plants have the potential to recruit similar nodular communities to native plants in greenhouse but not natural conditions (La Pierre et al., 2017), which would point to a root both in genetic compatibility of the partners and in competitive interactions in a mixed natural community.

### Natives and nonnatives differ in interactions with non-diazotrophic nodule members

The co-occurrence networks demonstrated more than 100 significant co-occurrences, the majority of which were positive. The diversity of interactors is striking andgenerally involve either two diazotrophs (likely in direct competition in the nodule environment) or a diazotroph and a fungus. Negative interactions between kingdoms most obviously arise from competition (e..g, direct rhizobia/mycorrhizal competition for photosynthate) but could also result from nutrient dynamics (alternative nitrogen or phosphorous limitation) at the plant level as well as source effects. However, a much larger set of fungi and non-diazotrophic bacteria were positively correlated with diazotrophic taxa, yielding candidates for mutualistic non-diazotrophic endophytic partners in nodule symbiosis (Muresu et al., 2008; Dudeja et al., 2012; De Meyer et al., 2015).

There was a prevalence of negative interactions in the co-occurrence network of non-native hosts (Figure 8B). Non-diazotrophs accounted for majority of the negative interactions in nonnative hosts while most of the negative associations in the native host network were attributed only to diazotrophs. Typically, the focus has been on how mutualisms result in positive impacts on invasion biology (Lu et al., 2016). However, (Pringle et al., 2009) discuss negative impacts of symbionts on competing species, which may facilitate the invasion process. For instance, invasions of *Alliaria* and *Centaurea* appear to be partly facilitated by disruptions these plants cause in mycorrhizal networks. Disruption of mycorrhizae is a possible explanation of our finding, as at least two of the negative interactors with *Rhizobium* (*Russula* and *Trametes*) are mycorrhizal plants. An important limitation of the co-occurrence analysis is that direct mutualistic participation in the symbiosis is only provisional even if the positive co-occurrence is significant and specific to diazotrophs that are the central players in the nodule environment.

**Figure 8.**
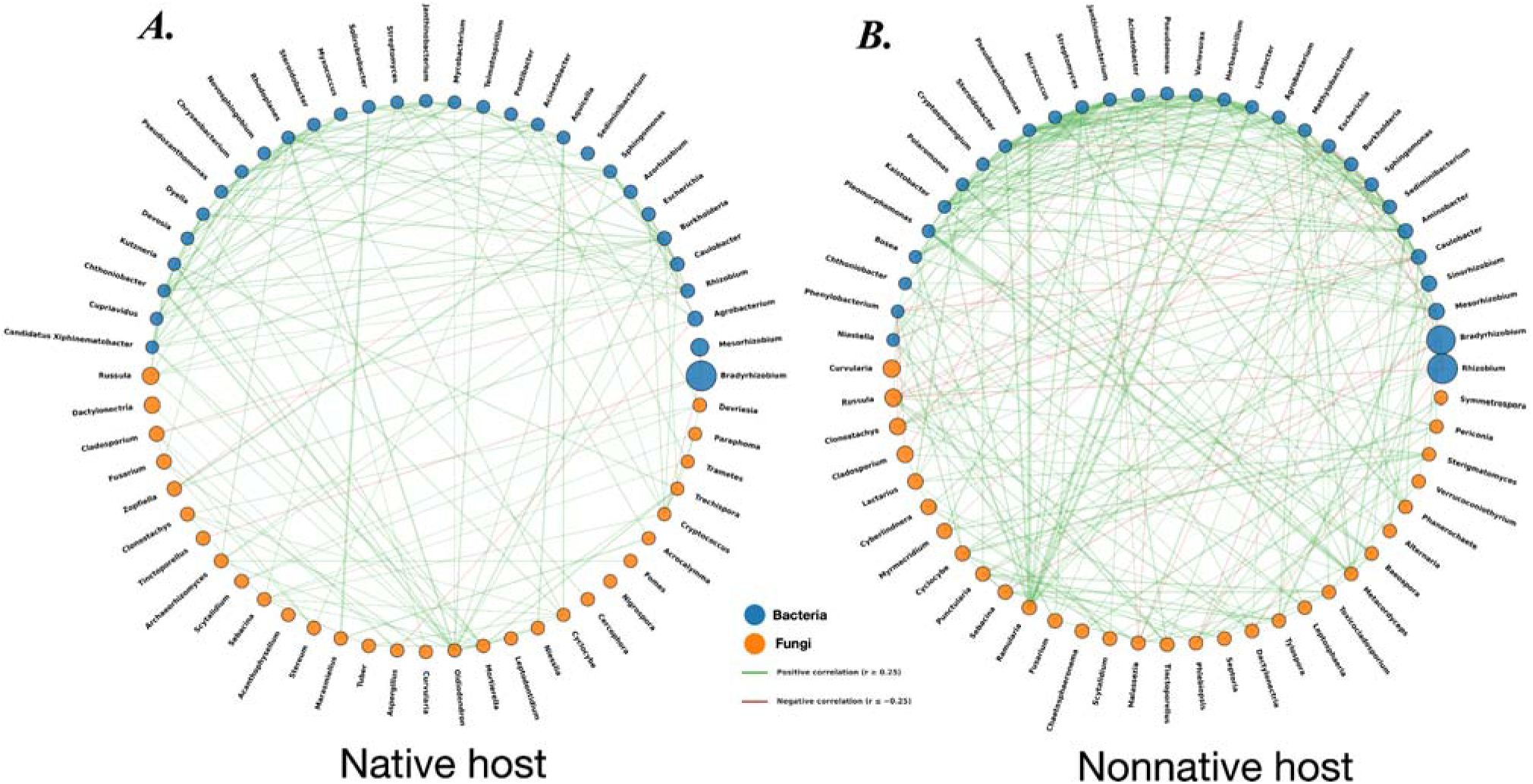
Bacterial-fungal co-occurrence network associated with (A) native (samples=71, nodes=60, edges=161) and (B) non-native (samples=36, nodes=60, edges=283) rhizobial legumes. Networks show the relationships among the 30 most abundant bacterial and fungal genera in native and non-native legume hosts. Nodes represent microbial genera, with blue representing bacteria and orange, fungi. Node size is proportional to the mean relative abundance of each genus. Edges indicate significant pairwise Spearman correlations in relative abundance (|r| ≥ 0.25), with green edges representing positive correlations and red edges representing negative correlations. Edge width is proportional to the strength of the correlation coefficient.

Alternative explanations such as correlated availability in species pools (possible if unlikely at the genus level) or correlated host plant interactions (e.g., negative interactions at the plant level with the plant then engaging in a specific diazotroph interaction, or two specific diazotrophic and mycorrhizal interactions mediated at the plant level) would need to be ruled out, likely in an experimental framework with these candidates in hand.

### Conclusions

We aimed to use a broad survey of nodulating plant microbiomes from the southeastern United States to investigate the influence of host plant phylogeny, habitat, and geographic origin to the diversity and composition of diazotrophic symbionts and their non-diazotrophic neighbors. We found evidence that host specificity appears to apply primarily at higher phylogenetic levels (“fuzzy specificity”) with looser host-symbiont taxonomic alignment at the species level. The effects seen for habitat and native origin were more nuanced and generally weaker. However, nonnative legumes, which represent a relatively narrow host phylogenetic range and largely disturbed habitats, exhibit significantly different symbiont composition from their neighboring native taxa. Overall, multiple factors shape nodule microbial communities with host phylogeny (specifically, membership in relatively divergent clades) displaying the strongest effect. Because diazotrophic transmission is strictly horizontal, phylogenetic effect likely represents conservatism of symbiotic function in the host rather than the direct effect of history. Future work should focus on adding larger ecological gradients which will be necessary to better investigate the role of abiotic environmental conditions on composition and diversity.

## Acknowledgments

This work was supported by the National Science Foundation (NSF DEB-2316266) and the Mississippi State University Strategic Research Initiative. Personnel from the Noxubee National Wildlife Refuge and the Tombigbee National Forest are thanked for permitting to collect.

## Author Contributions

D.J.P.: Conceptualization, formal analysis, visualization, writing–original draft, review, and editing. P.D.G.: analysis, visualization, writing–review and editing. J.D.: Fieldwork, writing–review and editing. S.A.: writing–review and editing. T.S.: laboratory work, writing–review and editing. N.J.E.W.: Fieldwork, laboratory work, writing–review and editing. C.M.S.: Fieldwork, laboratory work, writing–review and editing. H.R.J: laboratory protocol development, writing–review and editing. R.P.G.: writing–review and editing. R.A.F.: Conceptualization, fieldwork, laboratory work, analysis, writing–review and editing, funding acquisition, resources, supervision.

## Notes

### Competing Interest Statement

The authors have declared no competing interest.

### Summary of Updates

Added Tajinder Singh as a co-author. Added an author contributions section.

