## Supplement for "“Fuzzy specificity” shapes diazotroph diversity and composition in nodulating plants of the Southeastern USA"

### APPENDIX

**Appendix S1.** Provenance table for accessions included in this study.

| Sample ID | Host species | Host tribe | Symbiont | Sample type | Native? | Habitat | State | County | Latitude | Longitude |
|---|---|---|---|---|---|---|---|---|---|---|
| 1-1-E-No | *Stylosanthes biflora* | Dalbergieae | rhizobial | nodule | yes | pinewoods | MS | Winston | 33.214225 | -89.089824 |
| 1-1-E-Rh | *Stylosanthes biflora* | Dalbergieae | rhizobial | rhizosphere | yes | pinewoods | MS | Winston | 33.214225 | -89.089824 |
| 1-1-E-Ro | *Stylosanthes biflora* | Dalbergieae | rhizobial | root | yes | pinewoods | MS | Winston | 33.214225 | -89.089824 |
| 1-3-E-So | *Stylosanthes biflora* | Dalbergieae | rhizobial | soil | yes | pinewoods | MS | Winston | 33.214225 | -89.089824 |
| 10-1-E-So | *Elaeagnus pungens* | actinorhizal | actinorhizal | soil | no | streamside | MS | Oktibbeha | 33.459967 | -88.784658 |
| 2-1-E-No | *Desmodium* cf. *ciliare* | Desmodieae | rhizobial | nodule | yes | pinewoods | MS | Winston | 33.214225 | -89.089824 |
| 2-1-E-Rh | *Desmodium* cf. *ciliare* | Desmodieae | rhizobial | rhizosphere | yes | pinewoods | MS | Winston | 33.214225 | -89.089824 |
| 2-1-E-Ro | *Desmodium* cf. *ciliare* | Desmodieae | rhizobial | root | yes | pinewoods | MS | Winston | 33.214225 | -89.089824 |
| 2-1-E-So | *Desmodium* cf. *ciliare* | Desmodieae | rhizobial | soil | yes | pinewoods | MS | Winston | 33.214225 | -89.089824 |
| 29-D-No | *Albizia julbrissin* | Mimosoideae | rhizobial | nodule | no | uplandwoods | MS | Oktibbeha | 33.430673 | -88.766685 |
| 29-D-Ro | *Albizia julbrissin* | Mimosoideae | rhizobial | root | no | uplandwoods | MS | Oktibbeha | 33.430673 | -88.766685 |
| 29-E-No | *Albizia julbrissin* | Mimosoideae | rhizobial | nodule | no | uplandwoods | MS | Oktibbeha | 33.430673 | -88.766685 |
| 3-D-So | *Lespedeza bicolor* | Desmodieae | rhizobial | soil | no | uplandwoods | MS | Choctaw | 33.268855 | -89.140852 |
| 3-E-No | *Lespedeza bicolor* | Desmodieae | rhizobial | nodule | no | uplandwoods | MS | Choctaw | 33.268855 | -89.140852 |

| | | | | | | | | | | |
|---|---|---|---|---|---|---|---|---|---|---|
| 3-E-Rh | *Lespedeza bicolor* | Desmodieae | rhizobial | rhizosphere | no | uplandwoods | MS | Choctaw | 33.268855 | -89.140852 |
| 3-E-Ro | *Lespedeza bicolor* | Desmodieae | rhizobial | root | no | uplandwoods | MS | Choctaw | 33.268855 | -89.140852 |
| 4-E-No | *Strophostyles umbellata* | Phaseoleae | rhizobial | nodule | yes | uplandwoods | MS | Choctaw | 33.266809 | -89.13839 |
| 4-E-Rh | *Strophostyles umbellata* | Phaseoleae | rhizobial | rhizosphere | yes | uplandwoods | MS | Choctaw | 33.266809 | -89.13839 |
| 4-E-Ro | *Strophostyles umbellata* | Phaseoleae | rhizobial | root | yes | uplandwoods | MS | Choctaw | 33.266809 | -89.13839 |
| 4-E-So | *Strophostyles umbellata* | Phaseoleae | rhizobial | soil | yes | uplandwoods | MS | Choctaw | 33.266809 | -89.13839 |
| 5-E-No | *Hylodesmum glutinosum* | Desmodieae | rhizobial | nodule | yes | uplandwoods | MS | Choctaw | 33.268361 | -89.140016 |
| 5-E-Rh | *Hylodesmum glutinosum* | Desmodieae | rhizobial | rhizosphere | yes | uplandwoods | MS | Choctaw | 33.268361 | -89.140016 |
| 5-E-Ro | *Hylodesmum glutinosum* | Desmodieae | rhizobial | root | yes | uplandwoods | MS | Choctaw | 33.268361 | -89.140016 |
| 5-E-So | *Hylodesmum glutinosum* | Desmodieae | rhizobial | soil | yes | uplandwoods | MS | Choctaw | 33.268361 | -89.140016 |
| 6-2-E-No | *Sesbania herbacea* | Sesbanieae | rhizobial | nodule | yes | ditch | MS | Clay | 33.545891 | -88.632514 |
| 6-2-E-Rh | *Sesbania herbacea* | Sesbanieae | rhizobial | rhizosphere | yes | ditch | MS | Clay | 33.545891 | -88.632514 |
| 6-E-So | *Sesbania herbacea* | Sesbanieae | rhizobial | soil | yes | ditch | MS | Clay | 33.545891 | -88.632514 |
| 7-E-No | *Lespedeza cuneata* | Desmodieae | rhizobial | nodule | no | roadside | MS | Clay | 33.546661 | -88.632383 |
| 7-E-Rh | *Lespedeza cuneata* | Desmodieae | rhizobial | rhizosphere | no | roadside | MS | Clay | 33.546661 | -88.632383 |
| 7-E-Ro | *Lespedeza cuneata* | Desmodieae | rhizobial | root | no | roadside | MS | Clay | 33.546661 | -88.632383 |
| 7-E-So | *Lespedeza cuneata* | Desmodieae | rhizobial | soil | no | roadside | MS | Clay | 33.546661 | -88.632383 |

| | | | | | | | | | | |
|---|---|---|---|---|---|---|---|---|---|---|
| 8-1-E-No | *Robinia pseudoacacia* | Robinieae | rhizobial | nodule | yes | woodedge | MS | Oktibbeha | 33.459942 | -88.785164 |
| 8-1-E-Rh | *Robinia pseudoacacia* | Robinieae | rhizobial | rhizosphere | yes | woodedge | MS | Oktibbeha | 33.459942 | -88.785164 |
| 8-1-E-Ro | *Robinia pseudoacacia* | Robinieae | rhizobial | root | yes | woodedge | MS | Oktibbeha | 33.459942 | -88.785164 |
| 9-1-E-No | *Wisteria frutescens* | Millettieae | rhizobial | nodule | yes | streamside | MS | Oktibbeha | 33.294781 | -88.773088 |
| 9-1-E-Rh | *Wisteria frutescens* | Millettieae | rhizobial | rhizosphere | yes | streamside | MS | Oktibbeha | 33.294781 | -88.773088 |
| 9-1-E-Ro | *Wisteria frutescens* | Millettieae | rhizobial | root | yes | streamside | MS | Oktibbeha | 33.294781 | -88.773088 |
| 9-2-D-So | *Wisteria frutescens* | Millettieae | rhizobial | soil | yes | streamside | MS | Oktibbeha | 33.294781 | -88.773088 |
| CMS689-D-Rh | *Chamaecrista nictitans* | Cassieae | rhizobial | rhizosphere | yes | pinewoods | MS | Winston | 33.2149498 | -88.946612 |
| CMS689-E-No | *Chamaecrista nictitans* | Cassieae | rhizobial | nodule | yes | pinewoods | MS | Winston | 33.2149498 | -88.946612 |
| CMS690-D-Rh | *Lespedeza cuneata* | Desmodieae | rhizobial | rhizosphere | no | pinewoods | MS | Winston | 33.2149498 | -88.946612 |
| CMS690-E-Ro | *Lespedeza cuneata* | Desmodieae | rhizobial | root | no | pinewoods | MS | Winston | 33.2149498 | -88.946612 |
| CMS698-E-No | *Crotalaria sagittalis* | Crotalarieae | rhizobial | nodule | yes | pinewoods | MS | Choctaw | 33.242741 | -89.156933 |
| CMS698-E-Rh | *Crotalaria sagittalis* | Crotalarieae | rhizobial | rhizosphere | yes | pinewoods | MS | Choctaw | 33.242741 | -89.156933 |
| CMS698-E-Ro | *Crotalaria sagittalis* | Crotalarieae | rhizobial | root | yes | pinewoods | MS | Choctaw | 33.242741 | -89.156933 |
| CMS698-E-So | *Crotalaria sagittalis* | Crotalarieae | rhizobial | soil | yes | pinewoods | MS | Choctaw | 33.242741 | -89.156933 |
| JRD317-No | *Morella pumila* | actinorhizal | actinorhizal | nodule | yes | flatwoods | FL | Alachua | 29.7524 | -82.2184 |
| JRD321-2-E-No | *Desmodium incanum* | Desmodieae | rhizobial | nodule | no | ditch | FL | Alachua | 29.5889 | -82.3058 |

| | | | | | | | | | | |
|---|---|---|---|---|---|---|---|---|---|---|
| JRD321-No | *Desmodium incanum* | Desmodieae | rhizobial | nodule | no | ditch | FL | Alachua | 29.5889 | -82.3058 |
| JRD330-No | *Arachis hypogaea* | Dalbergieae | rhizobial | nodule | no | field | FL | Alachua | 29.6436 | -82.3549 |
| JRD331-No | *Desmodium incanum* | Desmodieae | rhizobial | nodule | no | field | FL | Alachua | 29.6436 | -82.3549 |
| JRD332-No | *Chamaecrista nictitans* | Cassieae | rhizobial | nodule | yes | woodedge | FL | Alachua | 29.6436 | -82.3549 |
| JRD335-No | *Crotalaria pallida* var. *obovata* | Crotalarieae | rhizobial | nodule | no | woodedge | FL | Alachua | 29.6436 | -82.3549 |
| JRD336-No | *Trifolium repens* | Trifolieae | rhizobial | nodule | no | field | FL | Alachua | 29.6436 | -82.3549 |
| JRD338-No | *Alysicarpus vaginalis* | Desmodieae | rhizobial | nodule | no | field | FL | Alachua | 29.6436 | -82.3549 |
| JRD348-No | *Crotalaria* sp. | Crotalarieae | rhizobial | nodule | na | woodedge | FL | Alachua | 29.6436 | -82.3549 |
| JRD349-No | *Crotalaria lanceolata* | Crotalarieae | rhizobial | nodule | no | woodedge | FL | Alachua | 29.6436 | -82.3549 |
| JRD386-2-E-No | *Mimosa* sp. | Mimosoideae | rhizobial | nodule | na | sandhill | FL | Alachua | 29.5889 | -82.3058 |
| JRD388-2-E-No | *Sesbania* sp. | Sesbanieae | rhizobial | nodule | yes | sandhill | FL | Alachua | 29.5889 | -82.3058 |
| JRD389-2-E-No | *Sesbania* sp. | Sesbanieae | rhizobial | nodule | yes | sandhill | FL | Alachua | 29.5889 | -82.3058 |
| JRD389-2-E-Ro | *Sesbania* sp. | Sesbanieae | rhizobial | root | yes | sandhill | FL | Alachua | 29.5889 | -82.3058 |
| JRD390-E-No | *Galactia regularis* | Diocleae | rhizobial | nodule | yes | sandhill | FL | Alachua | 29.5889 | -82.3058 |
| JRD392-E-No | *Desmodium strictum* | Desmodieae | rhizobial | nodule | yes | sandhill | FL | Alachua | 29.5889 | -82.3058 |
| JRD392-E-Rh | *Desmodium strictum* | Desmodieae | rhizobial | rhizosphere | yes | sandhill | FL | Alachua | 29.5889 | -82.3058 |
| JRD392-E-Ro | *Desmodium strictum* | Desmodieae | rhizobial | root | yes | sandhill | FL | Alachua | 29.5889 | -82.3058 |

| | | | | | | | | | | |
|---|---|---|---|---|---|---|---|---|---|---|
| JRD400-E-No | *Rynchosia* sp. | Phaseoleae | rhizobial | nodule | na | sandhill | FL | Alachua | 29.7294 | -82.4414 |
| JRD400-E-Rh | *Rynchosia* sp. | Phaseoleae | rhizobial | rhizosphere | na | sandhill | FL | Alachua | 29.7294 | -82.4414 |
| JRD400-E-Ro | *Rynchosia* sp. | Phaseoleae | rhizobial | root | na | sandhill | FL | Alachua | 29.7294 | -82.4414 |
| JRD405-E-No | *Dalea* sp. | Amorpheae | rhizobial | nodule | yes | sandhill | FL | Alachua | 29.7294 | -82.4414 |
| JRD405-E-Ro | *Dalea* sp. | Amorpheae | rhizobial | root | yes | sandhill | FL | Alachua | 29.7294 | -82.4414 |
| JRD419-E-No | *Crotalaria rotundifolia* | Crotalarieae | rhizobial | nodule | yes | flatwoods | FL | Alachua | 29.7524 | -82.2184 |
| JRD420-E-No | *Centrosema virginianum* | Phaseoleae | rhizobial | nodule | yes | flatwoods | FL | Alachua | 29.7524 | -82.2184 |
| JRD422-E-No | *Chamaecrista nictitans* | Cassieae | rhizobial | nodule | yes | flatwoods | FL | Alachua | 29.7524 | -82.2184 |
| JRD423-E-No | *Desmodium strictum* | Desmodieae | rhizobial | nodule | yes | flatwoods | FL | Alachua | 29.7524 | -82.2184 |
| JRD426-E-No | *Desmodium strictum* | Desmodieae | rhizobial | nodule | yes | flatwoods | FL | Alachua | 29.7524 | -82.2184 |
| JRD458-E-No | *Chamaecrista nictitans* | Cassieae | rhizobial | nodule | yes | sandhill | FL | Alachua | 29.7524 | -82.2184 |
| JRD461-E-No | *Lespedeza* sp. | Desmodieae | rhizobial | nodule | na | sandhill | FL | Alachua | 29.7524 | -82.2184 |
| JRD461-E-Rh | *Lespedeza* sp. | Desmodieae | rhizobial | rhizosphere | na | sandhill | FL | Alachua | 29.7524 | -82.2184 |
| JRD462-E-No | *Lespedeza* sp. | Desmodieae | rhizobial | nodule | na | sandhill | FL | Alachua | 29.7524 | -82.2184 |
| JRD462-E-Rh | *Lespedeza* sp. | Desmodieae | rhizobial | rhizosphere | na | sandhill | FL | Alachua | 29.7524 | -82.2184 |
| JRD474-E-No | *Chamaecrista nictitans* | Cassieae | rhizobial | nodule | yes | sandhill | FL | Jefferson | 30.4312 | -83.8897 |
| JRD476-E-No | *Clitoria mariana* | Phaseoleae | rhizobial | nodule | yes | sandhill | FL | Jefferson | 30.4312 | -83.8897 |

| | | | | | | | | | | |
|---|---|---|---|---|---|---|---|---|---|---|
| JRD477-E-No | *Clitoria mariana* | Phaseoleae | rhizobial | nodule | yes | sandhill | FL | Jefferson | 30.4312 | -83.8897 |
| JRD477-E-Ro | *Clitoria mariana* | Phaseoleae | rhizobial | root | yes | sandhill | FL | Jefferson | 30.4312 | -83.8897 |
| JRD478-E-No | *Fabaceae* sp. | na | rhizobial | nodule | na | sandhill | FL | Jefferson | 30.4312 | -83.8897 |
| JRD478-E-Ro | *Fabaceae* sp. | na | rhizobial | root | na | sandhill | FL | Jefferson | 30.4312 | -83.8897 |
| JRD480-E-No | *Desmodium floridanum* | Desmodieae | rhizobial | nodule | yes | sandhill | FL | Jefferson | 30.4312 | -83.8897 |
| JRD484-E-No | *Tephrosia* sp. | Millettieae | rhizobial | nodule | na | sandhill | FL | Jefferson | 30.4312 | -83.8897 |
| JRD485-E-No | *Desmodium floridanum* | Desmodieae | rhizobial | nodule | yes | sandhill | FL | Jefferson | 30.4312 | -83.8897 |
| JRD485-E-Ro | *Desmodium floridanum* | Desmodieae | rhizobial | root | yes | sandhill | FL | Jefferson | 30.4312 | -83.8897 |
| JRD487-E-No | *Lespedeza* sp. | Desmodieae | rhizobial | nodule | na | sandhill | FL | Jefferson | 30.4312 | -83.8897 |
| JRD489-E-No | *Desmodium rotundifolium* | Desmodieae | rhizobial | nodule | yes | sandhill | FL | Jefferson | 30.4312 | -83.8897 |
| JRD489-E-Ro | *Desmodium rotundifolium* | Desmodieae | rhizobial | root | yes | sandhill | FL | Jefferson | 30.4312 | -83.8897 |
| JRD494-E-No | *Lespedeza cuneata* | Desmodieae | rhizobial | nodule | no | woodedge | AL | Cleburne | 33.7504 | -85.6247 |
| JRD496-E-No | *Amphicarpaea bracteata* | Phaseoleae | rhizobial | nodule | yes | woodedge | AL | Cleburne | 33.7504 | -85.6247 |
| JRD498-E-No | *Desmodium* sp. | Desmodieae | rhizobial | nodule | na | woodedge | AL | Cleburne | 33.7504 | -85.6247 |
| JRD499-E-No | *Chamaecrista nictitans* | Cassieae | rhizobial | nodule | yes | woodedge | AL | Cleburne | 33.7504 | -85.6247 |
| JRD501-E-No | *Centrosema virginianum* | Phaseoleae | rhizobial | nodule | yes | woodedge | AL | Cleburne | 33.7504 | -85.6247 |
| JRD511-E-No | *Desmodium strictum* | Desmodieae | rhizobial | nodule | yes | pinewoods | AL | Cleburne | 33.7504 | -85.6247 |

| | | | | | | | | | | |
|---|---|---|---|---|---|---|---|---|---|---|
| JRD512-E-No | *Desmodium strictum* | Desmodieae | rhizobial | nodule | yes | pinewoods | AL | Cleburne | 33.7504 | -85.6247 |
| JRD514-E-No | *Desmodium floridanum* | Desmodieae | rhizobial | nodule | yes | pinewoods | AL | Cleburne | 33.7504 | -85.6247 |
| JRD525-E-No | Fabaceae sp. | n/a | rhizobial | nodule | na | pinewoods | AL | Cleburne | 33.7504 | -85.6247 |
| N1-3-E-No | *Vicia caroliniana* | Fabeae | rhizobial | nodule | yes | uplandwoods | MS | Winston | 33.208977 | -89.074675 |
| N1-3-E-Ro | *Vicia caroliniana* | Fabeae | rhizobial | root | yes | uplandwoods | MS | Winston | 33.208977 | -89.074675 |
| N10-1-E-No | *Trifolium resupinatum* | Trifolieae | rhizobial | nodule | no | roadside | MS | Oktibbeha | 33.44731 | -88.697626 |
| N10-4-D-No | *Trifolium resupinatum* | Trifolieae | rhizobial | nodule | no | roadside | MS | Oktibbeha | 33.44731 | -88.697626 |
| N11-1-E-No | *Trifolium campestre* | Trifolieae | rhizobial | nodule | no | field | MS | Oktibbeha | 33.542046 | -88.77493 |
| N11-1-E-Rh | *Trifolium campestre* | Trifolieae | rhizobial | rhizosphere | no | field | MS | Oktibbeha | 33.542046 | -88.77493 |
| N11-2-D-No | *Trifolium campestre* | Trifolieae | rhizobial | nodule | no | field | MS | Oktibbeha | 33.542046 | -88.77493 |
| N11-2-D-Rh | *Trifolium campestre* | Trifolieae | rhizobial | rhizosphere | no | field | MS | Oktibbeha | 33.542046 | -88.77493 |
| N12-1-E-No | *Trifolium dubium* | Trifolieae | rhizobial | nodule | no | field | MS | Oktibbeha | 33.542046 | -88.77493 |
| N12-1-E-Rh | *Trifolium dubium* | Trifolieae | rhizobial | rhizosphere | no | field | MS | Oktibbeha | 33.542046 | -88.77493 |
| N12-4-D-No | *Trifolium dubium* | Trifolieae | rhizobial | nodule | no | field | MS | Oktibbeha | 33.542046 | -88.77493 |
| N12-4-D-Rh | *Trifolium dubium* | Trifolieae | rhizobial | rhizosphere | no | field | MS | Oktibbeha | 33.542046 | -88.77493 |
| N13-2-D-No | *Trifolium lappaceum* | Trifolieae | rhizobial | nodule | no | roadside | MS | Oktibbeha | 33.54222 | -88.77489 |
| N13-2-D-Rh | *Trifolium lappaceum* | Trifolieae | rhizobial | rhizosphere | no | roadside | MS | Oktibbeha | 33.54222 | -88.77489 |

| | | | | | | | | | | |
|---|---|---|---|---|---|---|---|---|---|---|
| N13-3-E-No | *Trifolium lappaceum* | Trifolieae | rhizobial | nodule | no | roadside | MS | Oktibbeha | 33.54222 | -88.77489 |
| N13-3-E-Rh | *Trifolium lappaceum* | Trifolieae | rhizobial | rhizosphere | no | roadside | MS | Oktibbeha | 33.54222 | -88.77489 |
| N13-3-E-Ro | *Trifolium lappaceum* | Trifolieae | rhizobial | root | no | roadside | MS | Oktibbeha | 33.54222 | -88.77489 |
| N14-3-E-No | *Orbexilum pedunculatum* | Psoraleeae | rhizobial | nodule | yes | field | MS | Oktibbeha | 33.381723 | -88.962821 |
| N14-4-D-No | *Orbexilum pedunculatum* | Psoraleeae | rhizobial | nodule | yes | field | MS | Oktibbeha | 33.381723 | -88.962821 |
| N15-3-E-No | *Ceanothus americanus* | actinorhizal | actinorhizal | nodule | yes | woodedge | MS | Choctaw | 33.277697 | -89.14036 |
| N16-1-E-No | *Alnus serratula* | actinorhizal | actinorhizal | nodule | yes | streamside | MS | Choctaw | 33.276387 | -89.137866 |
| N16-5-D-No | *Alnus serratula* | actinorhizal | actinorhizal | nodule | yes | streamside | MS | Choctaw | 33.276387 | -89.137866 |
| N17-8-D-No | *Elaeagnus pungens* | actinorhizal | actinorhizal | nodule | yes | streamside | MS | Oktibbeha | 33.459967 | -88.784658 |
| N17-E-No | *Elaeagnus pungens* | actinorhizal | actinorhizal | nodule | yes | streamside | MS | Oktibbeha | 33.459967 | -88.784658 |
| N18-2-D-No | *Vicia villosa* | Fabeae | rhizobial | nodule | no | field | MS | Oktibbeha | 33.403166 | -88.847269 |
| N18-5-E-No | *Vicia villosa* | Fabeae | rhizobial | nodule | no | field | MS | Oktibbeha | 33.403166 | -88.847269 |
| N18-5-E-Rh | *Vicia villosa* | Fabeae | rhizobial | rhizosphere | no | field | MS | Oktibbeha | 33.403166 | -88.847269 |
| N18-5-E-Ro | *Vicia villosa* | Fabeae | rhizobial | root | no | field | MS | Oktibbeha | 33.403166 | -88.847269 |
| N19-3-E-No | *Amorpha fruticosa* | Amorpheae | rhizobial | nodule | yes | bottomlandw oods | MS | Oktibbeha | 33.360551 | -88.721584 |
| N19-D-So | *Amorpha fruticosa* | Amorpheae | rhizobial | soil | yes | bottomlandw oods | MS | Oktibbeha | 33.360551 | -88.721584 |
| N2-1-E-No | *Vicia grandiflora* | Fabeae | rhizobial | nodule | no | field | MS | Winston | 33.18157 | -89.01218 |

| | | | | | | | | | | |
|---|---|---|---|---|---|---|---|---|---|---|
| N2-1-E-Ro | *Vicia grandiflora* | Fabeae | rhizobial | root | no | field | MS | Winston | 33.18157 | -89.01218 |
| N20-1-E-No | *Morella cerifera* | actinorhizal | actinorhizal | nodule | yes | bottomlandw oods | MS | Winston | 33.277964 | -88.900427 |
| N20-8-D-No | *Morella cerifera* | actinorhizal | actinorhizal | nodule | yes | bottomlandw oods | MS | Winston | 33.277964 | -88.900427 |
| N21-1-D-No | *Trifolium arvense* | Trifolieae | rhizobial | nodule | no | field | MS | Winston | 33.199285 | -88.994276 |
| N21-1-D-Ro | *Trifolium arvense* | Trifolieae | rhizobial | root | no | field | MS | Winston | 33.199285 | -88.994276 |
| N21-3-E-No | *Trifolium arvense* | Trifolieae | rhizobial | nodule | no | field | MS | Winston | 33.199285 | -88.994276 |
| N21-3-E-Rh | *Trifolium arvense* | Trifolieae | rhizobial | rhizosphere | no | field | MS | Winston | 33.199285 | -88.994276 |
| N21-D-So | *Trifolium arvense* | Trifolieae | rhizobial | soil | no | field | MS | Winston | 33.199285 | -88.994276 |
| N22-1-E-No | *Tephrosia virginica* | Millettieae | rhizobial | nodule | yes | pinewoods | MS | Winston | 33.264949 | -89.090208 |
| N22-1-E-Rh | *Tephrosia virginica* | Millettieae | rhizobial | rhizosphere | yes | pinewoods | MS | Winston | 33.264949 | -89.090208 |
| N23-2-D-No | *Securigera varia* | Loteae | rhizobial | nodule | no | field | MS | Oktibbeha | 33.468642 | -88.778935 |
| N23-3-E-No | *Securigera varia* | Loteae | rhizobial | nodule | no | field | MS | Oktibbeha | 33.468642 | -88.778935 |
| N23-3-E-Rh | *Securigera varia* | Loteae | rhizobial | rhizosphere | no | field | MS | Oktibbeha | 33.468642 | -88.778935 |
| N24-3-E-No | *Wisteria sinensis* | Millettieae | rhizobial | nodule | no | roadside | MS | Winston | 33.234035 | -89.065799 |
| N24-3-E-Ro | *Wisteria sinensis* | Millettieae | rhizobial | root | no | roadside | MS | Winston | 33.234035 | -89.065799 |
| N24-4-D-No | *Wisteria sinensis* | Millettieae | rhizobial | nodule | no | roadside | MS | Winston | 33.234035 | -89.065799 |
| N24-4-D-Ro | *Wisteria sinensis* | Millettieae | rhizobial | root | no | roadside | MS | Winston | 33.234035 | -89.065799 |

| | | | | | | | | | | |
|---|---|---|---|---|---|---|---|---|---|---|
| N25-2-D-No | *Pueraria montana* | Glycininae | rhizobial | nodule | no | roadside | MS | Winston | 33.234035 | -89.065799 |
| N25-5-E-No | *Pueraria montana* | Glycininae | rhizobial | nodule | no | roadside | MS | Winston | 33.234035 | -89.065799 |
| N25-5-E-Rh | *Pueraria montana* | Glycininae | rhizobial | rhizosphere | no | roadside | MS | Winston | 33.234035 | -89.065799 |
| N26-2-D-No | *Wisteria frutescens* | Millettieae | rhizobial | nodule | yes | streamside | MS | Clay | 33.661639 | -88.512679 |
| N26-2-D-Ro | *Wisteria frutescens* | Millettieae | rhizobial | root | yes | streamside | MS | Clay | 33.661639 | -88.512679 |
| N26-3-E-No | *Wisteria frutescens* | Millettieae | rhizobial | nodule | yes | streamside | MS | Clay | 33.661639 | -88.512679 |
| N26-3-E-Rh | *Wisteria frutescens* | Millettieae | rhizobial | rhizosphere | yes | streamside | MS | Clay | 33.661639 | -88.512679 |
| N26-3-E-Ro | *Wisteria frutescens* | Millettieae | rhizobial | root | yes | streamside | MS | Clay | 33.661639 | -88.512679 |
| N27-1-E-No | *Tephrosia onobrychoides* | Millettieae | rhizobial | nodule | yes | pinewoods | MS | Oktibbeha | 33.315172 | -88.839268 |
| N27-2-E-No | *Tephrosia onobrychoides* | Millettieae | rhizobial | nodule | yes | pinewoods | MS | Oktibbeha | 33.315172 | -88.839268 |
| N28-1-E-No | *Desmanthus illinoensis* | Mimosoideae | rhizobial | nodule | yes | prairie | MS | Monroe | 33.80743 | -88.508842 |
| N28-2-D-No | *Desmanthus illinoensis* | Mimosoideae | rhizobial | nodule | yes | prairie | MS | Monroe | 33.80743 | -88.508842 |
| N29-1-E-No | *Crotalaria spectabilis* | Crotalarieae | rhizobial | nodule | no | streamside | MS | Monroe | 33.806818 | -88.511614 |
| N29-1-E-Rh | *Crotalaria spectabilis* | Crotalarieae | rhizobial | rhizosphere | no | streamside | MS | Monroe | 33.806818 | -88.511614 |
| N29-6-D-No | *Crotalaria spectabilis* | Crotalarieae | rhizobial | nodule | no | streamside | MS | Monroe | 33.806818 | -88.511614 |
| N3-3-E-No | *Lathyrus hirsutus* | Fabeae | rhizobial | nodule | no | field | MS | Oktibbeha | 33.484168 | -88.796163 |
| N30-2-E-No | *Dalea purpurea* | Amorpheae | rhizobial | nodule | yes | calcareousprairie | MS | Oktibbeha | 33.507114 | -88.737653 |

| | | | | | | | | | | |
|---|---|---|---|---|---|---|---|---|---|---|
| N30-2-E-Rh | *Dalea purpurea* | Amorpheae | rhizobial | rhizosphere | yes | calcareouspra irie | MS | Oktibbeha | 33.507114 | -88.737653 |
| N30-3-E-No | *Dalea purpurea* | Amorpheae | rhizobial | nodule | yes | calcareouspra irie | MS | Oktibbeha | 33.507114 | -88.737653 |
| N30-3-E-Ro | *Dalea purpurea* | Amorpheae | rhizobial | root | yes | calcareouspra irie | MS | Oktibbeha | 33.507114 | -88.737653 |
| N31-1-E-No | *Neptunia lutea* | Mimosoideae | rhizobial | nodule | yes | calcareouspra irie | MS | Oktibbeha | 33.507128 | -88.737673 |
| N31-1-E-Rh | *Neptunia lutea* | Mimosoideae | rhizobial | rhizosphere | yes | calcareouspra irie | MS | Oktibbeha | 33.507128 | -88.737673 |
| N31-3-E-No | *Neptunia lutea* | Mimosoideae | rhizobial | nodule | yes | calcareouspra irie | MS | Oktibbeha | 33.507128 | -88.737673 |
| N32-7-E-No | *Hylodesmum pauciflorum* | Desmodieae | rhizobial | nodule | yes | uplandwoods | MS | Oktibbeha | 33.424227 | -88.76036 |
| N32-8-D-No | *Hylodesmum pauciflorum* | Desmodieae | rhizobial | nodule | yes | uplandwoods | MS | Oktibbeha | 33.424227 | -88.76036 |
| N32-8-D-Ro | *Hylodesmum pauciflorum* | Desmodieae | rhizobial | root | yes | uplandwoods | MS | Oktibbeha | 33.424227 | -88.76036 |
| N33-1-E-No | *Galactia regularis* | Diocleae | rhizobial | nodule | yes | calcareouspra irie | MS | Oktibbeha | 33.506564 | -88.736927 |
| N33-1-E-Ro | *Galactia regularis* | Diocleae | rhizobial | root | yes | calcareouspra irie | MS | Oktibbeha | 33.506564 | -88.736927 |
| N33-2-D-No | *Galactia regularis* | Diocleae | rhizobial | nodule | yes | calcareouspra irie | MS | Oktibbeha | 33.506564 | -88.736927 |
| N33-2-D-Ro | *Galactia regularis* | Diocleae | rhizobial | root | yes | calcareouspra irie | MS | Oktibbeha | 33.506564 | -88.736927 |
| N34-1-E-No | *Desmodium sessilifolium* | Desmodieae | rhizobial | nodule | yes | calcareouspra irie | MS | Oktibbeha | 33.506668 | -88.736796 |
| N34-2-D-No | *Desmodium sessilifolium* | Desmodieae | rhizobial | nodule | yes | calcareouspra irie | MS | Oktibbeha | 33.506668 | -88.736796 |
| N34-2-D-Ro | *Desmodium sessilifolium* | Desmodieae | rhizobial | root | yes | calcareouspra irie | MS | Oktibbeha | 33.506668 | -88.736796 |
| N35-4-D-No | *Sesbania herbacea* | Sesbanieae | rhizobial | nodule | yes | ditch | MS | Oktibbeha | 33.466646 | -88.786337 |

| | | | | | | | | | | |
|---|---|---|---|---|---|---|---|---|---|---|
| N35-4-D-Ro | *Sesbania herbacea* | Sesbanieae | rhizobial | root | yes | ditch | MS | Oktibbeha | 33.466646 | -88.786337 |
| N35-5-E-No | *Sesbania herbacea* | Sesbanieae | rhizobial | nodule | yes | ditch | MS | Oktibbeha | 33.466646 | -88.786337 |
| N36-3-E-No | *Sesbania vesicaria* | Sesbanieae | rhizobial | nodule | yes | streamside | MS | Clay | 33.637192 | -88.501584 |
| N36-3-E-Rh | *Sesbania vesicaria* | Sesbanieae | rhizobial | rhizosphere | yes | streamside | MS | Clay | 33.637192 | -88.501584 |
| N36-4-D-No | *Sesbania herbacea* | Sesbanieae | rhizobial | nodule | yes | ditch | MS | Clay | 33.637192 | -88.501584 |
| N36-4-D-Ro | *Sesbania herbacea* | Sesbanieae | rhizobial | root | yes | ditch | MS | Clay | 33.637192 | -88.501584 |
| N36-D-So | *Sesbania herbacea* | Sesbanieae | rhizobial | soil | yes | ditch | MS | Clay | 33.637192 | -88.501584 |
| N37-1-E-No | *Centrosema virginianum* | Phaseoleae | rhizobial | nodule | yes | woodedge | MS | Clay | 33.637577 | -88.503273 |
| N37-1-E-Ro | *Centrosema virginianum* | Phaseoleae | rhizobial | root | yes | woodedge | MS | Clay | 33.637577 | -88.503273 |
| N37-2-E-No | *Centrosema virginianum* | Phaseoleae | rhizobial | nodule | yes | woodedge | MS | Clay | 33.637577 | -88.503273 |
| N37-2-E-Ro | *Centrosema virginianum* | Phaseoleae | rhizobial | root | yes | woodedge | MS | Clay | 33.637577 | -88.503273 |
| N38-1-E-No | *Aeschynomene indica* | Dalbergieae | rhizobial | nodule | no | streamside | MS | Lowndes | 33.52539 | -88.468494 |
| N38-1-E-Rh | *Aeschynomene indica* | Dalbergieae | rhizobial | rhizosphere | no | streamside | MS | Lowndes | 33.52539 | -88.468494 |
| N38-1-E-Ro | *Aeschynomene indica* | Dalbergieae | rhizobial | root | no | streamside | MS | Lowndes | 33.52539 | -88.468494 |
| N38-2-D-No | *Aeschynomene indica* | Dalbergieae | rhizobial | nodule | no | streamside | MS | Lowndes | 33.52539 | -88.468494 |
| N38-2-D-Ro | *Aeschynomene indica* | Dalbergieae | rhizobial | root | no | streamside | MS | Lowndes | 33.52539 | -88.468494 |
| N39-2-D-No | *Apios americana* | Phaseoleae | rhizobial | nodule | yes | streamside | MS | Lowndes | 33.525403 | -88.468489 |

| | | | | | | | | | | |
|---|---|---|---|---|---|---|---|---|---|---|
| N39-2-D-Ro | *Apios americana* | Phaseoleae | rhizobial | root | yes | streamside | MS | Lowndes | 33.525403 | -88.468489 |
| N39-3-E-No | *Apios americana* | Phaseoleae | rhizobial | nodule | yes | streamside | MS | Lowndes | 33.525403 | -88.468489 |
| N39-3-E-Rh | *Apios americana* | Phaseoleae | rhizobial | rhizosphere | yes | streamside | MS | Lowndes | 33.525403 | -88.468489 |
| N4-D-No | *Vicia tetrasperma* | Fabeae | rhizobial | nodule | no | field | MS | Oktibbeha | 33.506432 | -88.736572 |
| N4-D-So | *Vicia tetrasperma* | Fabeae | rhizobial | soil | no | field | MS | Oktibbeha | 33.506432 | -88.736572 |
| N40-2-D-No | *Kummerowia striata* | Desmodieae | rhizobial | nodule | no | streamside | MS | Lowndes | 33.525328 | -88.468254 |
| N40-2-D-Ro | *Kummerowia striata* | Desmodieae | rhizobial | root | no | streamside | MS | Lowndes | 33.525328 | -88.468254 |
| N40-3-E-No | *Kummerowia striata* | Desmodieae | rhizobial | nodule | no | streamside | MS | Lowndes | 33.525328 | -88.468254 |
| N40-3-E-Rh | *Kummerowia striata* | Desmodieae | rhizobial | rhizosphere | no | streamside | MS | Lowndes | 33.525328 | -88.468254 |
| N40-3-E-Ro | *Kummerowia striata* | Desmodieae | rhizobial | root | no | streamside | MS | Lowndes | 33.525328 | -88.468254 |
| N41-1-E-No | *Mimosa microphylla* | Mimosoideae | rhizobial | nodule | yes | pinewoods | MS | Oktibbeha | 33.315501 | -88.838927 |
| N41-1-E-Rh | *Mimosa microphylla* | Mimosoideae | rhizobial | rhizosphere | yes | pinewoods | MS | Oktibbeha | 33.315501 | -88.838927 |
| N41-2-D-No | *Mimosa microphylla* | Mimosoideae | rhizobial | nodule | yes | pinewoods | MS | Oktibbeha | 33.315501 | -88.838927 |
| N41-2-D-Ro | *Mimosa microphylla* | Mimosoideae | rhizobial | root | yes | pinewoods | MS | Oktibbeha | 33.315501 | -88.838927 |
| N42-3-E-No | *Amphicarpaea bracteata* | Phaseoleae | rhizobial | nodule | yes | woodedge | MS | Oktibbeha | 33.425293 | -88.759451 |
| N42-D-4-No | *Amphicarpaea bracteata* | Phaseoleae | rhizobial | nodule | yes | woodedge | MS | Oktibbeha | 33.425293 | -88.759451 |
| N42-D-4-Ro | *Amphicarpaea bracteata* | Phaseoleae | rhizobial | root | yes | woodedge | MS | Oktibbeha | 33.425293 | -88.759451 |

| | | | | | | | | | | |
|---|---|---|---|---|---|---|---|---|---|---|
| N43-3-E-No | *Desmodium ciliare* | Desmodieae | rhizobial | nodule | yes | pinewoods | MS | Oktibbeha | 33.315615 | -88.846867 |
| N43-3-E-Ro | *Desmodium ciliare* | Desmodieae | rhizobial | root | yes | pinewoods | MS | Oktibbeha | 33.315615 | -88.846867 |
| N43-D-4-No | *Desmodium ciliare* | Desmodieae | rhizobial | nodule | yes | pinewoods | MS | Oktibbeha | 33.315615 | -88.846867 |
| N43-D-So | *Desmodium ciliare* | Desmodieae | rhizobial | soil | yes | pinewoods | MS | Oktibbeha | 33.315615 | -88.846867 |
| N44-2-D-Ro | *Lespedeza virginica* | Desmodieae | rhizobial | root | yes | pinewoods | MS | Oktibbeha | 33.315621 | -88.846871 |
| N44-3-E-No | *Lespedeza virginica* | Desmodieae | rhizobial | nodule | yes | pinewoods | MS | Oktibbeha | 33.315621 | -88.846871 |
| N44-3-E-Ro | *Lespedeza virginica* | Desmodieae | rhizobial | root | yes | pinewoods | MS | Oktibbeha | 33.315621 | -88.846871 |
| N45-3-E-No | *Desmodium rotundifolium* | Desmodieae | rhizobial | nodule | yes | uplandwoods | MS | Oktibbeha | 33.4237 | -88.759645 |
| N45-D-4-No | *Desmodium rotundifolium* | Desmodieae | rhizobial | nodule | yes | uplandwoods | MS | Oktibbeha | 33.4237 | -88.759645 |
| N45-D-4-Ro | *Desmodium rotundifolium* | Desmodieae | rhizobial | root | yes | uplandwoods | MS | Oktibbeha | 33.4237 | -88.759645 |
| N46-5-E-Rh | *Senna obtusifolia* | Cassieae | rhizobial | rhizosphere | no | pinewoods | MS | Oktibbeha | 33.315913 | -88.850061 |
| N46-D-8-Ro | *Senna obtusifolia* | Cassieae | rhizobial | root | no | pinewoods | MS | Oktibbeha | 33.315913 | -88.850061 |
| N47-3-E-No | *Desmodium paniculatum* | Desmodieae | rhizobial | nodule | yes | pinewoods | MS | Oktibbeha | 33.315798 | -88.849432 |
| N47-D-2-No | *Desmodium paniculatum* | Desmodieae | rhizobial | nodule | yes | pinewoods | MS | Oktibbeha | 33.315798 | -88.849432 |
| N48-3-E-No | *Desmodium glabellum* | Desmodieae | rhizobial | nodule | yes | pinewoods | MS | Oktibbeha | 33.315501 | -88.838927 |
| N48-D-4-No | *Desmodium glabellum* | Desmodieae | rhizobial | nodule | yes | pinewoods | MS | Oktibbeha | 33.315501 | -88.838927 |
| N48-D-4-Ro | *Desmodium glabellum* | Desmodieae | rhizobial | root | yes | pinewoods | MS | Oktibbeha | 33.315501 | -88.838927 |

| | | | | | | | | | | |
|---|---|---|---|---|---|---|---|---|---|---|
| N49-1-E-No | *Lackeya multiflora* | Diocleae | rhizobial | nodule | yes | bottomlandwoods | MS | Oktibbeha | 33.28885 | -88.772152 |
| N49-1-E-Rh | *Lackeya multiflora* | Diocleae | rhizobial | rhizosphere | yes | bottomlandwoods | MS | Oktibbeha | 33.28885 | -88.772152 |
| N49-1-E-Ro | *Lackeya multiflora* | Diocleae | rhizobial | root | yes | bottomlandwoods | MS | Oktibbeha | 33.28885 | -88.772152 |
| N49-3-D-No | *Lackeya multiflora* | Diocleae | rhizobial | nodule | yes | bottomlandwoods | MS | Oktibbeha | 33.28885 | -88.772152 |
| N49-3-D-Ro | *Lackeya multiflora* | Diocleae | rhizobial | root | yes | bottomlandwoods | MS | Oktibbeha | 33.28885 | -88.772152 |
| N5-4-D-No | *Trifolium incarnatum* | Trifolieae | rhizobial | nodule | no | field | MS | Oktibbeha | 33.484218 | -88.795596 |
| N50-5-E-No | *Strophostyles helvola* | Phaseoleae | rhizobial | nodule | yes | streamside | MS | Clay | 33.637009 | -88.501852 |
| N50-5-E-Ro | *Strophostyles helvola* | Phaseoleae | rhizobial | root | yes | streamside | MS | Clay | 33.637009 | -88.501852 |
| N50-D-So | *Strophostyles helvola* | Phaseoleae | rhizobial | soil | yes | streamside | MS | Clay | 33.637009 | -88.501852 |
| N51-1-E-No | *Chamaecrista fasciculata* | Cassieae | rhizobial | nodule | yes | field | MS | Clay | 33.636251 | -88.524455 |
| N51-2-D-No | *Chamaecrista fasciculata* | Cassieae | rhizobial | nodule | yes | field | MS | Clay | 33.636251 | -88.524455 |
| N6-1-E-No | *Vicia sativa* | Fabeae | rhizobial | nodule | no | field | MS | Oktibbeha | 33.542731 | -88.773684 |
| N6-2-D-No | *Vicia sativa* | Fabeae | rhizobial | nodule | no | field | MS | Oktibbeha | 33.542731 | -88.773684 |
| N6-2-D-Ro | *Vicia sativa* | Fabeae | rhizobial | root | no | field | MS | Oktibbeha | 33.542731 | -88.773684 |
| N7-2-D-No | *Melilotus alba* | Trifolieae | rhizobial | nodule | no | calcareousprairie | MS | Oktibbeha | 33.507366 | -88.737845 |
| N7-2-D-Ro | *Melilotus alba* | Trifolieae | rhizobial | root | no | calcareousprairie | MS | Oktibbeha | 33.507366 | -88.737845 |
| N7-3-E-No | *Melilotus alba* | Trifolieae | rhizobial | nodule | no | calcareousprairie | MS | Oktibbeha | 33.507366 | -88.737845 |

| | | | | | | | | | | |
|---|---|---|---|---|---|---|---|---|---|---|
| N7-3-E-Ro | *Melilotus alba* | Trifolieae | rhizobial | root | no | calcareousprairie | MS | Oktibbeha | 33.507366 | -88.737845 |
| N8-1-E-No | *Trifolium pratense* | Trifolieae | rhizobial | nodule | no | field | MS | Oktibbeha | 33.484157 | -88.795428 |
| N8-1-E-Rh | *Trifolium pratense* | Trifolieae | rhizobial | rhizosphere | no | field | MS | Oktibbeha | 33.484157 | -88.795428 |
| N8-4-D-No | *Trifolium pratense* | Trifolieae | rhizobial | nodule | no | field | MS | Oktibbeha | 33.484157 | -88.795428 |
| N9-1-E-No | *Baptisia alba* | Sophoreae | rhizobial | nodule | yes | field | MS | Oktibbeha | 33.403215 | -88.839992 |
| N9-1-E-Rh | *Baptisia alba* | Sophoreae | rhizobial | rhizosphere | yes | field | MS | Oktibbeha | 33.403215 | -88.839992 |
| N9-1-E-Ro | *Baptisia alba* | Sophoreae | rhizobial | root | yes | field | MS | Oktibbeha | 33.403215 | -88.839992 |
| N9-4-D-No | *Baptisia alba* | Sophoreae | rhizobial | nodule | yes | field | MS | Oktibbeha | 33.403215 | -88.839992 |
| N9-D-So | *Baptisia alba* | Sophoreae | rhizobial | soil | yes | field | MS | Oktibbeha | 33.403215 | -88.839992 |

**Appendix S2.** Shannon diversity alpha rarefaction curves for bacterial 16S based on 1000 sequence rarefaction analyses from QIIME 2. Note: We used 100 for nodule, root, and rhizosphere, and 200 for soil as sequence cutoffs for downstream diversity analyses.

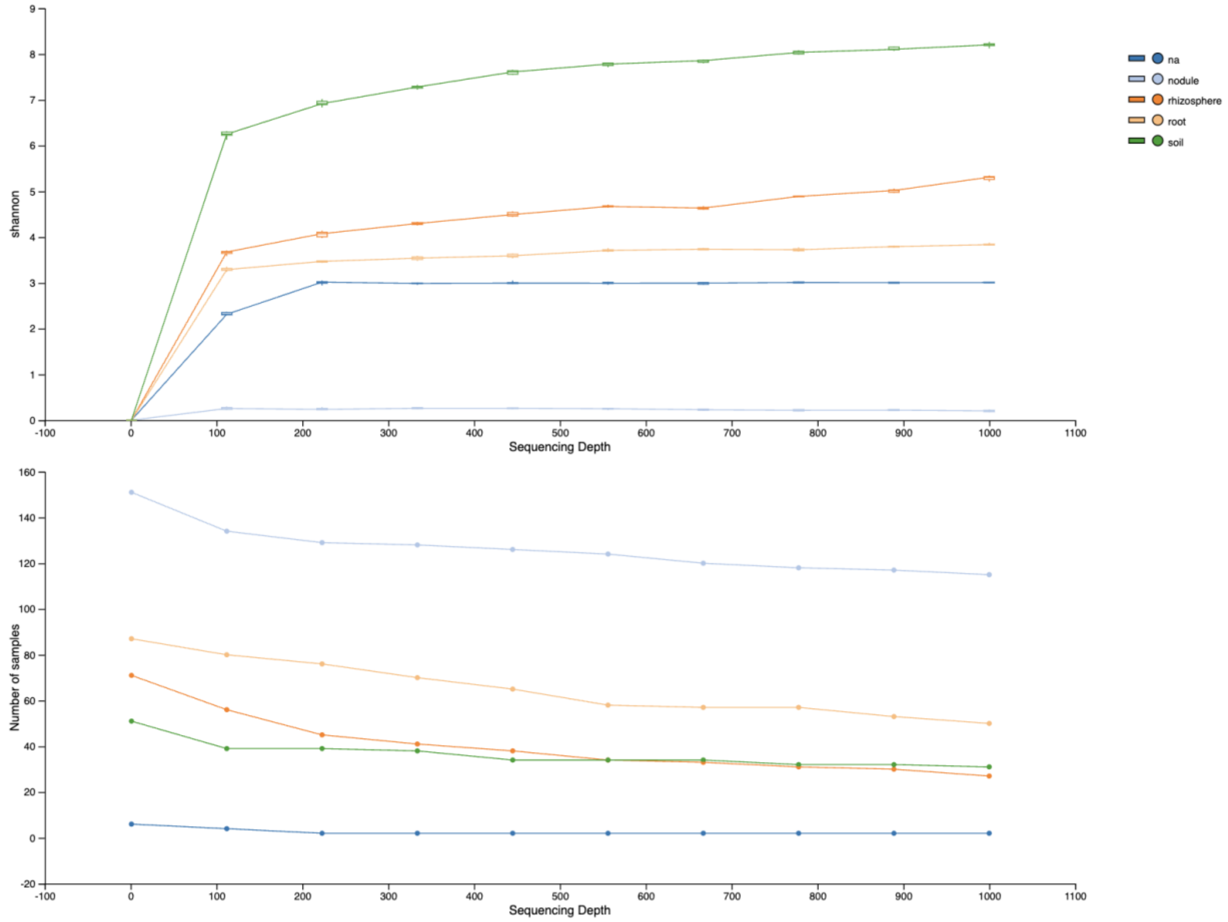


**Appendix S3.** A. Faith's Phylogenetic (top panel) and Shannon diversity (bottom panel) of nodular bacterial communities across nodulating host plant species. The two diversity metrics were not significantly different at the host species level.

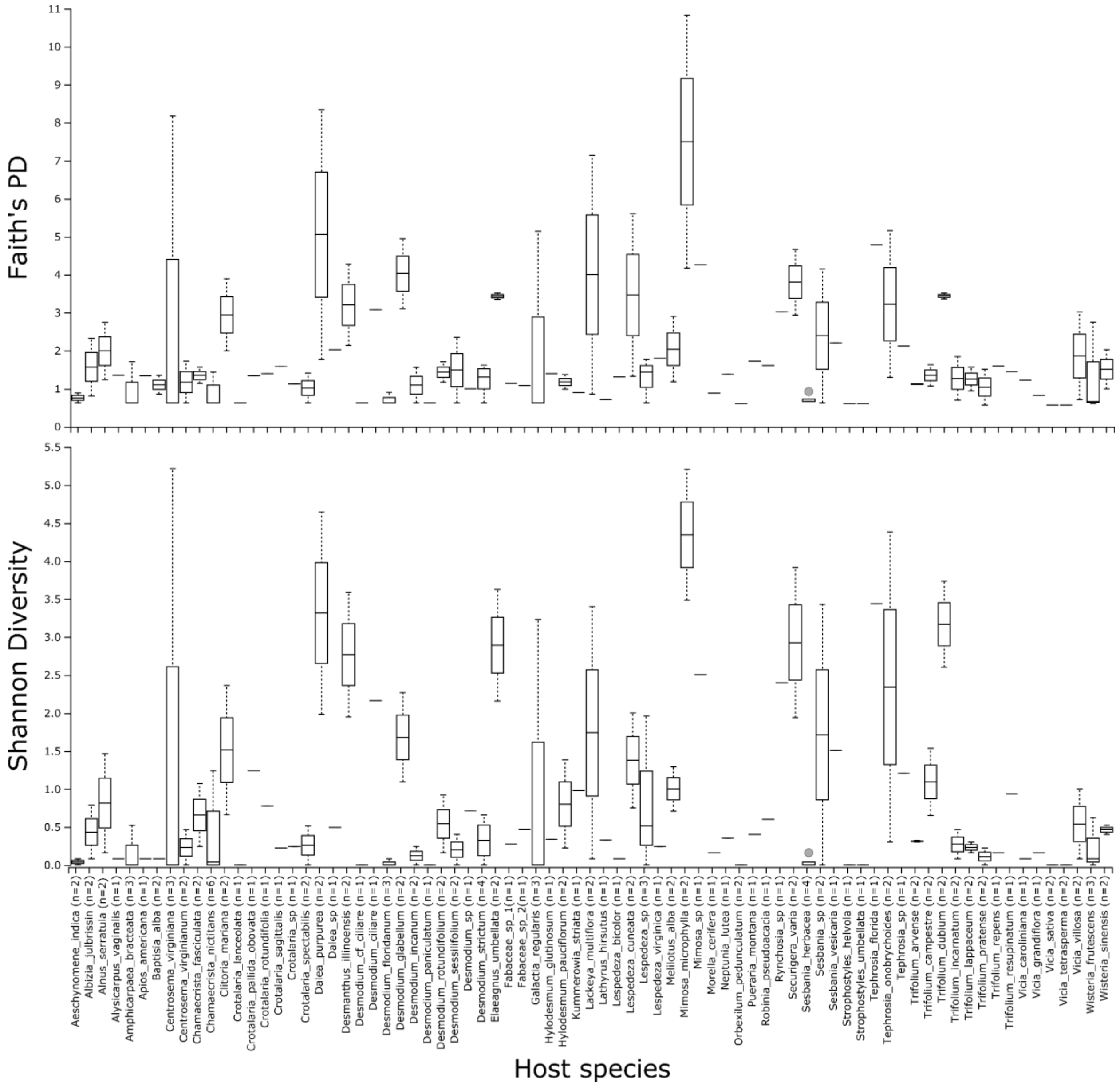


**Appendix S4.** A. Faith's Phylogenetic (top panel) and Shannon diversity (bottom panel) of nodular bacterial communities across nodulating host plant genera. The two diversity metrics were significantly different at the host genus level.

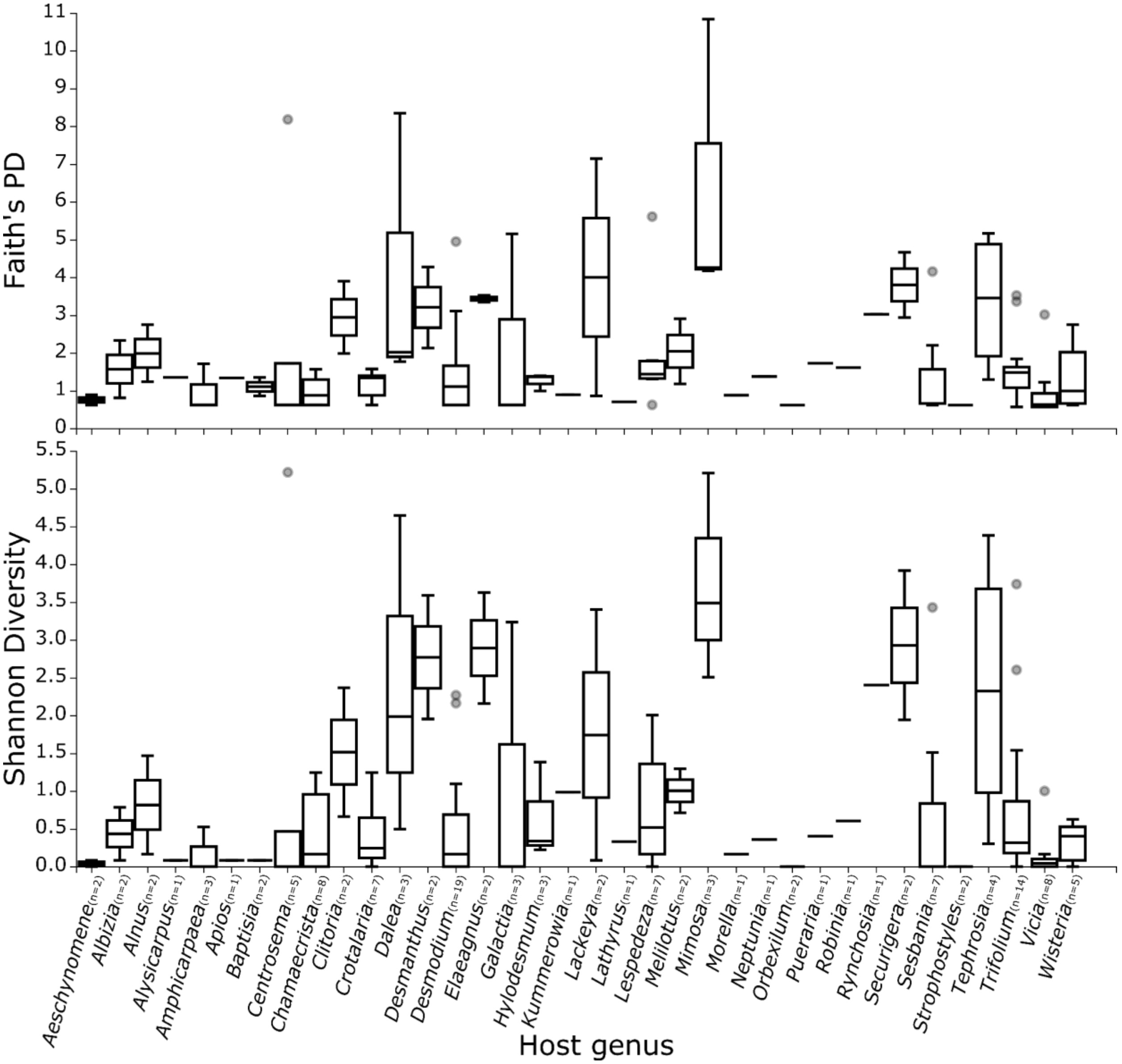


**Appendix S5.** A. Level 6 (genus) taxa barplots showing relative frequencies of bacteria between native and nonnative host plants across sampling levels (soil, rhizosphere, and root). B. ANCOM

Volcano plots across sampling levels showing no significant features that are differentially abundant between native and nonnative host plants.

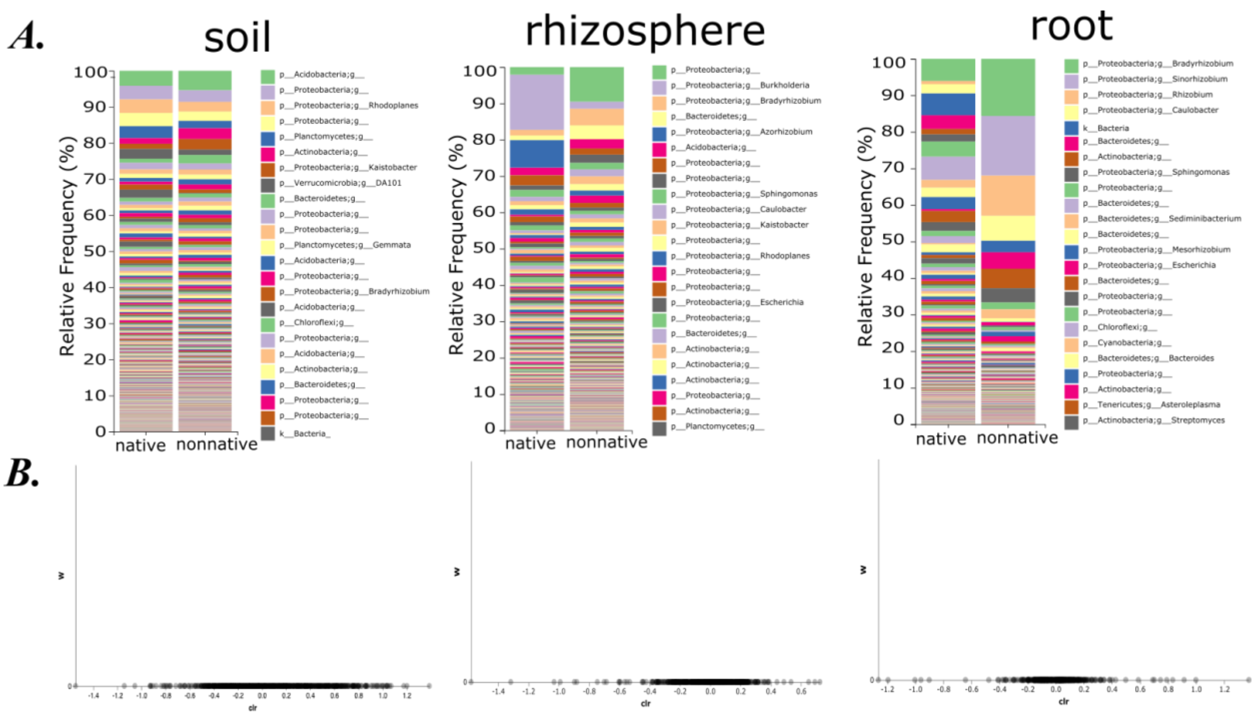

**Appendix S6.** Results of Mantel correlation test between nodule bacterial community UniFrac distance and host phylogenetic distance.

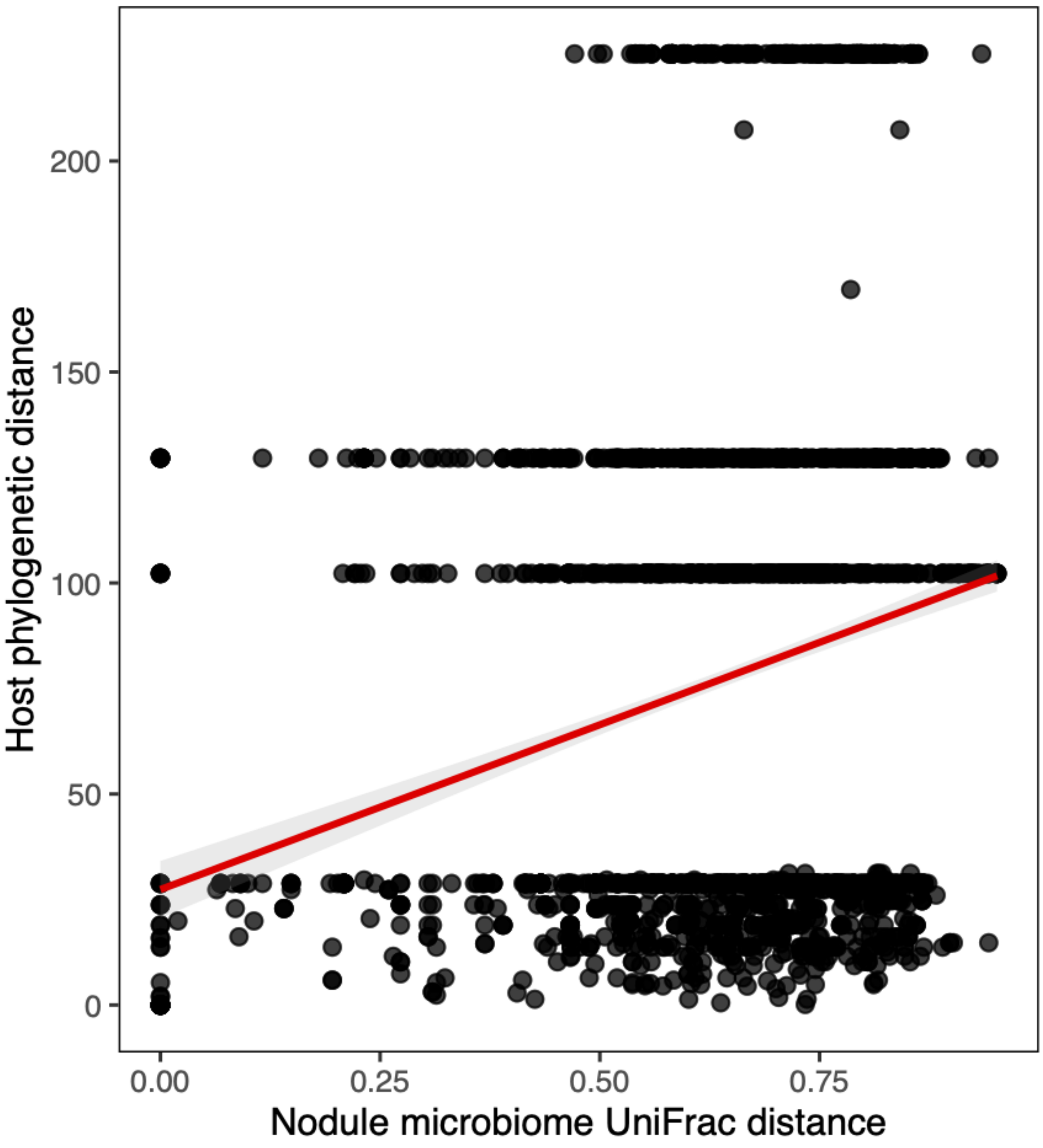


**Appendix S7.** Nodule Extraction Protocol

Use filter tips for all steps and work under a laminar flow/PCR hood unless otherwise noted. Stock solutions are reported at the end.

#### Sample preparation

##### Soil (in laminar flow hood):

**Materials (sample prep + Day one):** Get Eppendorf 2 mL round-bottom tubes (order number 022363352, 2*n* + 2 where *n* is the number of samples).

1. Dessicated and frozen soil should be set aside; these are ready to transfer to the lysis buffer.
2. For ethanol-preserved soil, spin down for 2 minutes, and pour off the ethanol onto a paper towel.
3. **Add “So” to the soil sample tube labels.**
4. Proceed to Extraction procedure: Day one.

##### Root samples (in the laminar flow hood):

**Materials and prep (Root samples + Day one):**

Get Eppendorf 2 mL round-bottom tubes (6*n* + 2 where *n* is the number of samples), 4 small beakers, sterile petri dishes (one per sample), clean tweezers and blades, HPLC water, 70% EtOH, 5% bleach.

Label 2 mL round-bottom tubes with the original sample name (1 per sample).

Label 2 mL round-bottom tubes adding Ro (2 per sample), No (2 per sample), Rh (1 per sample) to the original sample name.

1. Decontaminate the laminar flow hood with 50% bleach, check to remove any bleach drops on the walls, and run under UV for 15 minutes. Label tubes during the wait if not pre-labelled in advance.
2. If dry or frozen, add molecular grade water to ~top of the tube, vigorously flick, and wait 5 to 15 minutes; **if ethanol-preserved, proceed to (3)**.
3. Vigorously flick (this is the second time for dry and frozen materials).
4. Remove the root/nodule sample to a second identically labeled clean tube, **reserving the eluate** as the **rhizosphere sample, and adding “Rh” to the sample label**.
    a. If small root pieces remain in the rhizosphere sample, remove and discard.
5. For the rhizosphere sample, spin down 5 minutes at 10,000 g, then pour off the supernatant, reserving the pellet.
6. In the second tube with the root/nodule sample, add 70% ethanol (EtOH) to ~top of the tube, incubate for 1 minute, then pour off ethanol onto a paper towel. **(Skip for ethanol-preserved samples)**
7. Add 5% bleach to ~top of the tube, incubate for 1 minute, then pour off the bleach onto paper towel.

8. **Immediately** add molecular-grade water to ~top of tube, then pour off the water onto a paper towel.
9. Add molecular-grade water a second time to ~top of the tube, then pour off the water onto paper towel.
10. Move the root sample onto a clean petri dish; dissect out nodules from non-nodular root and transfer to a third identically labelled tube as the **nodule sample, adding "No" to the sample label**.
11. Return the roots without nodules to the second tube as the **root endophyte sample, adding "Ro" to the sample label**.
12. Proceed to the Extraction procedure: Day one.

**Extraction procedure: Day One**

**Materials and prep:**

Get Eppendorf 2 mL round-bottom tubes (*n* + 2), sterile beads, lysis buffer stock, potassium acetate stock.

1. Decontaminate the laminar flow hood if not immediately following dissection and sample preparation.
2. Prepare 2 sets of extra tubes for positive and negative controls and complete all the following steps for them along with the samples. Turn on the water bath and set it to 65° C for later.
3. Add 4-6 sterile metal beads to each tube.
4. Add 500 µL of lysis buffer to each tube. Additionally, add 500 µL of killed *E. coli* to the **positive control tube only**.
5. Process all tubes using the bead mill program (reported in Methods). Make sure to balance each of the 4 sectors of the bead beater and close securely without misthreading.
6. If large chunks of tissue are visible, process in the bead beater for a second time using program 2.
7. Ensure the tubes are closed after processing with the bead beater. Check the writing on the tubes and relabel if needed.
8. Incubate 20 minutes in a 65° C water bath. Meanwhile, add 200 µL of potassium acetate to a new set of pre-labelled tubes for later.
9. Check the writing on the tubes and relabel them if needed.
10. Process again with the bead beater using program 2.
11. Ensure tubes are closed and incubate for 10 minutes in a 65°C water bath.
12. Centrifuge 2 minutes at 10,000 g.
13. Transfer the supernatant to the pre-prepared 2 mL potassium acetate tube, take care to avoid the pellet and precipitated material as much as possible.
14. Chill in -20° C freezer overnight.
    a. Can also proceed after 1 hour to Day Two, but it is more efficient to work with larger sample numbers over multiple days.

#### Extraction procedure: Day Two

**Materials and prep:**

Get Eppendorf 2 mL round-bottom tubes (2*n* where *n* is the number of samples), spin filter columns (*n*), guanidine hydrochloride stock, wash solution stock, 80% EtOH, elution solution.

1. Centrifuge potassium acetate tubes with sample from Day One for 30 min at 10,000 g. Meanwhile, repeat the laminar flow hood decontamination procedure as in Day One and label all the tubes required for the protocol.
2. Prepare a new set of 2 mL tubes and add 1.2 mL guanidine hydrochloride (600 µL twice).
3. After the samples are done centrifuging, transfer the supernatant to pre-prepared labeled guanidine hydrochloride tubes and mix them using the pipette while adding, take care to avoid the pellet and precipitated material as much as possible.
4. Transfer 700 µL to a labelled spin filter.
5. Centrifuge for 2 minutes at 10,000 g.
6. **Discard flow-through, keeping the spin filter**.
7. Repeat steps 4 to 6 until you have transferred the entire volume through the spin filter, taking care to not cross-contaminate samples.
8. Add 500 µL wash solution.
9. Spin for 2 minutes at 10,000 g.
10. **Discard flow-through, keeping the spin filter**.
11. Repeat steps 8, 9, and 10 for a second wash.
12. Add 500 µL 80% ethanol (EtOH).
13. Spin for 2 minutes at 10,000 g.
14. **Discard flow-through, keeping the spin filter**.
15. Spin for 5 minutes at 10,000 g to ensure the spin column is dry.
16. Transfer the spin column to a new labeled tube (discard the old tube it was sitting in).
17. Add 200 µL elution solution for amplicon applications; add 50 µL elution solution for library preparation for shotgun sequencing and similar applications.
18. Incubate for 10 minutes at room temperature.
19. Spin for 2 minutes at 10,000 g.
20. Discard the spin filter, **keeping the eluate**.
21. Quantify via Qubit.
22. Store at -20°C.

#### Stock Solutions

Lysis solution (for 500 mL):

50 mL 1 *M* Tris pH 8.0
50 mL 0.5 *M* EDTA
50 mL 0.5 *M* NaCl (or 5 mL 5 *M* NaCl)
6.5 g SDS

343.5 mL $H_2O$ (if using 5 *M* NaCl, 388.5 mL $H_2O$)

Then autoclave.
When cool, add 100 mg RNAse A.

Potassium acetate (for 250 mL):
122.7 g potassium acetate
127.3 mL $H_2O$

Guanidine hydrochloride (for 500 mL):
31.5 g guanidine hydrochloride
316.5 mL ethanol
152 mL $H_2O$

Wash solution (for 1 L):
10 mL 1 *M* Tris pH 8.0
2 mL 0.5 *M* EDTA
10 mL 0.5 *M* NaCl (or 1 mL 5 *M* NaCl)
670 mL absolute ethanol
308 mL $H_2O$ (if using 5 *M* NaCl, 317 mL $H_2O$)

Elution solution (for 500 mL):
5 mL 1 *M* Tris pH 8.0
495 mL $H_2O$
